# Deschloroclozapine, a potent and selective chemogenetic actuator enables rapid neuronal and behavioral modulations in mice and monkeys

**DOI:** 10.1101/854513

**Authors:** Yuji Nagai, Naohisa Miyakawa, Hiroyuki Takuwa, Yukiko Hori, Kei Oyama, Bin Ji, Manami Takahashi, Xi-Ping Huang, Samuel T. Slocum, Jeffrey F. DiBerto, Yan Xiong, Takuya Urushihata, Toshiyuki Hirabayashi, Atsushi Fujimoto, Koki Mimura, Justin G. English, Jing Liu, Ken-ichi Inoue, Katsushi Kumata, Chie Seki, Maiko Ono, Masafumi Shimojo, Ming-Rong Zhang, Yutaka Tomita, Jin Nakahara, Tetsuya Suhara, Masahiko Takada, Makoto Higuchi, Jian Jin, Bryan L. Roth, Takafumi Minamimoto

**Affiliations:** Department of Functional Brain Imaging, National Institute of Radiological Sciences, National Institutes for Quantum and Radiological Science and Technology, Chiba Japan; Department of Pharmacology, University of North Carolina at Chapel Hill School of Medicine, Chapel Hill, NC, U.S.A.; Mount Sinai Center for Therapeutics Discovery, Departments of Pharmacological Sciences and Oncological Sciences, Tisch Cancer Institute, Icahn School of Medicine at Mount Sinai, New York, NY 10029, USA; Systems Neuroscience Section, Primate Research Institute, Kyoto University, Inuyama, Aichi, Japan; PRESTO, Japan Science and Technology Agency, Kawaguchi, Saitama, Japan; Department of Radiopharmaceuticals Development, National Institute of Radiological Sciences, National Institutes for Quantum and Radiological Science and Technology, Chiba, Japan; Department of Neurology, Keio University School of Medicine, Tokyo, Japan; Division of Chemical Biology and Medicinal Chemistry, Eshelman School of Pharmacy, University of North Carolina at Chapel Hill, Chapel Hill, NC, USA; National Institute of Mental Health Psychoactive Drug Screening Program (NIMH PDSP), Department of Pharmacology, University of North Carolina at Chapel Hill Medical School, Chapel Hill, NC, USA

## Abstract

The chemogenetic technology Designer Receptors Exclusively Activated by Designer Drugs (DREADDs) affords remotely reversible control of cellular signaling, neuronal activity and behavior. Although the combination of muscarinic-based DREADDs with clozapine-N-oxide (CNO) has been widely used, sluggish kinetics, metabolic liabilities, and potential off-target effects of CNO represent areas for improvement. Here we provide a new high affinity and selective agonist deschloroclozapine (DCZ) for muscarinic-based DREADDs. Positron emission tomography revealed that DCZ selectively bound to and occupied DREADDs in both mice and monkeys. Systemic delivery of low doses of DCZ (1 or 3 μg/kg) enhanced neuronal activity via hM_3_Dq within minutes in mice and monkeys. Intramuscular injections of DCZ (100 μg/kg) reversibly induced spatial working memory deficits in monkeys expressing hM_4_Di in the prefrontal cortex. DCZ represents the most potent, selective, metabolically stable and fast-acting DREADD agonist reported with utility in both mice and non-human primates for a variety of applications.

## Introduction

The chemogenetic technology Designer Receptors Exclusively Activated by Designer Drugs (DREADDs) affords a minimally invasive means to reversibly and remotely control the activity of DREADD-expressing cells by systemic delivery of DREADD ligands^1^. Several DREADDs exist, derived from muscarinic or kappa-opioid receptors^1,2^, and DREADDs have been widely adopted for neuroscience research^3^. Muscarinic receptor DREADDs are the most widely used and can be activated by clozapine-N-oxide (CNO) (Fig. 1d, right). Activation of hM_3_Dq enhances activity, while hM_4_Di silences neuronal activity. CNO activated DREADDs have been successfully applied in a variety of *in vitro* and *in vivo* contexts, including non-human primate studies to modify neural network activity^4^ and behavior^5–7^.

**Fig. 1.**
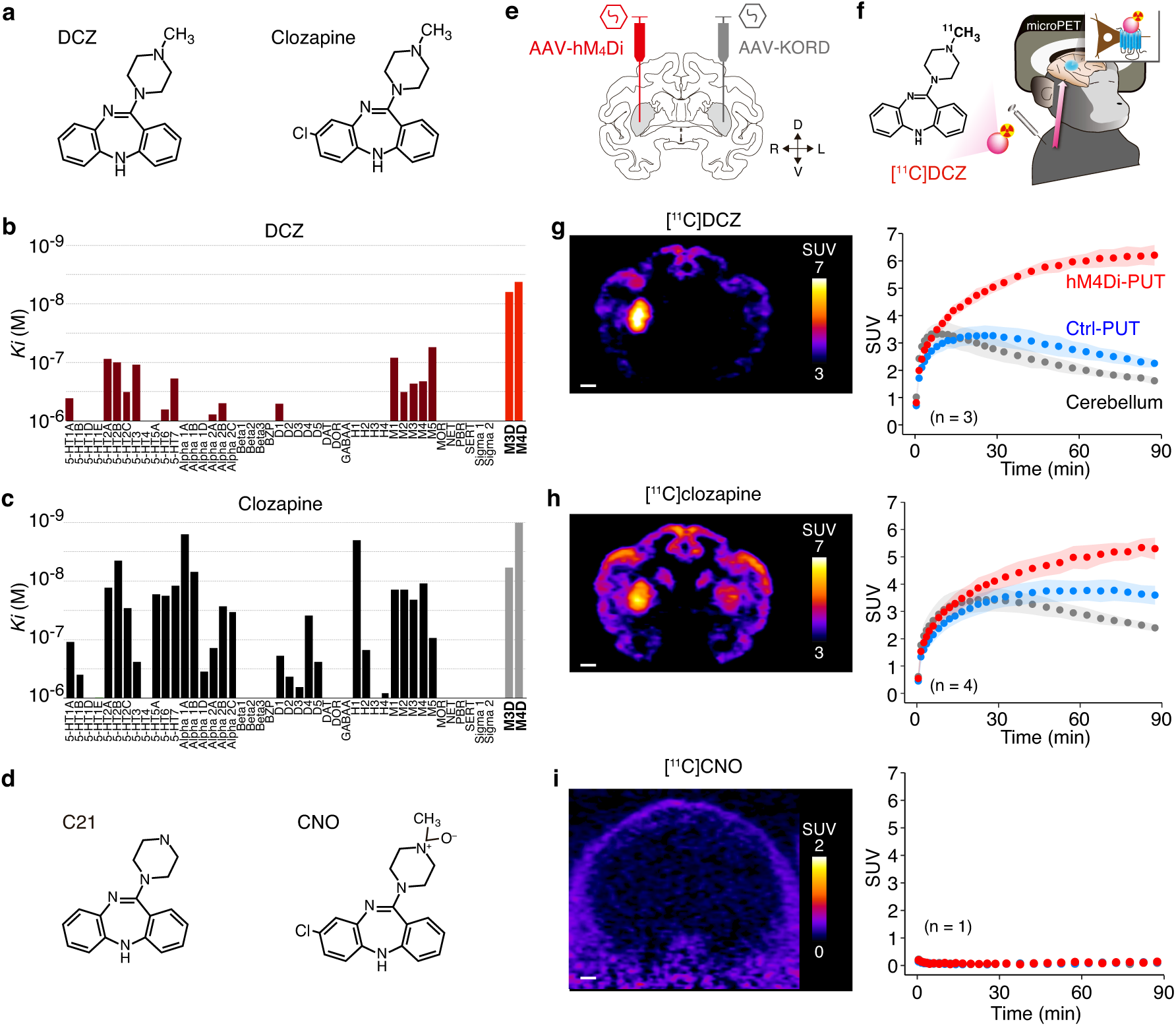
DCZ selectively binds to DREADDs. **a**, Chemical structure of DCZ and clozapine. **b**,**c**, Binding affinities of DCZ and clozapine to DREADDs and endogenous receptors, channels and transporters. *Ki* values are the average of at least 3 triplicate experiments with standard deviation values that are 3-fold less than the average. The values are also shown in Table 1 and Supplementary Table 1. **d**, Chemical structure of C21 and CNO. **e**, Illustration representing location of viral vector injections. AAV2-CMV-hM_4_Di and AAV2-CMV-KORD (as control) were injected into the right and left putamen, respectively. **f**, Illustration representing *in vivo* PET imaging with [^11^C]DCZ in monkeys. **g-i**, Representative PET imaging data obtained from monkey #209. **g**, (left) Coronal section of PET image with [^11^C]DCZ representing standardized radio-ligand uptake value [SUV; regional radioactivity (Bq/cm^3^) × body weight (g) / injected radioactivity (Bq)] between 30-90 min from injection. (right) Time course of regional uptake (mean ± sd, n = 3 scans) of [^11^C]DCZ at the hM_4_Di-expression region in the putamen (hM_4_Di-PUT), control region in the contralateral putamen (Ctrl-PUT), and the cerebellum, respectively. **h**, Same as **g** but with [^11^C]clozapine (n = 4). **i**, Same as **g** but with [^11^C]CNO (n = 1). Scale bars represent 5 mm.

Since CNO has modest brain permeability, relatively large systemic doses may be required for DREADD activation. Moreover, CNO can be metabolized to clozapine (Fig. 1a, right)—a potent, brain permeable DREADD agonist^1,8^. CNO metabolism to clozapine is significant in rodents^8,9^ and in monkeys^10^. In addition to activating DREADDs, clozapine potently binds to many endogenous receptors and transporters^11^; moreover, CNO or clozapine can produce confounding off-target side effects. Although alternative DREADD agonists, such as compound 21 (C21) (Fig. 1d, left) and perlapine^12,13^, have been recently developed, they require relatively large systemic doses to activate DREADDs *in vivo*, and may have off-target actions^13^. Given the broad utility and popularity of muscarinic-based DREADDs, the development of a selective, high-affinity, metabolically stable, and brain-penetrable DREADD agonist is important.

**Table 1.** Binding affinities of DCZ, clozapine and CNO to DREADDs and native muscarinic receptors.

| compound | *K<sub>i</sub>* (nM) |  |  |  |
| --- | --- | --- | --- | --- |
|  | *pK<sub>i</sub>* (± SEM) |  |  |  |
|  | hM<sub>3</sub>Dq | hM<sub>3</sub> | hM<sub>4</sub>Di | hM<sub>4</sub> |
| DCZ | 6.3 | 230 | 4.2 | 210 |
|  | 8.20 (± 0.15) | 6.63 (± 0.06) | 8.38 (± 0.27) | 6.68 (± 0.04) |
| clozapine | 5.9 | 21 | 0.89 | 10 |
|  | 8.23 (± 0.26) | 7.69 (± 0.14) | 9.05 (± 0.13) | 7.98 (± 0.10) |
| CNO | 680 | 2600 | 360 | 3700 |
|  | 6.17 (± 0.07) | 5.59 (± 0.06) | 6.44 (± 0.03) | 5.43 (± 0.18) |
| C21 | 850 | 270 | 178* | 3,631* |
|  | 6.07 (± 0.06) | 6.57 (± 0.15) | 6.75 (± 0.26) | 5.44 (± 0.11) |

Mean *K<sub>i</sub>* values (upper row) and mean ± SEM of *pK<sub>i</sub>* values (lower row) are shown. The average of at least 3 duplicate experiments with standard deviation (SD) values that are 3-fold less than average. \* data from Thompson et al.<sup>12</sup>. Radioligand binding assays were performed with [<sup>3</sup>H]quinuclidinyl benzilate (QNB) on HEK293T cells. The average [<sup>3</sup>H]QNB concentrations were 0.8 nM for hM<sub>3</sub>/hM<sub>4</sub> and 2.0 nM for hM<sub>3</sub>Dq/hM<sub>4</sub>Di.

Here we identified a suitable DREADD agonist that we have named DCZ [deschloroclozapine or 11-(4-methyl-1-piperazinyl)-5H-dibenzo(b,e)(1,4)diazepine] (Fig. 1a, left), derived from our previous studies on DREADD ligands for muscarinic-based DREADDs^12^. We became aware of DCZ as it had been previously reported to have substantially lower affinity for dopaminergic (D_1_ and D_2_) and serotonergic (5-HT_2A_ and 5-HT_2C_) receptors than clozapine^14^. We determined that DCZ has 100-fold improved affinity and greater agonist potency for hM_3_Dq and hM_4_Di relative to CNO or C21, with reduced off-target binding compared with clozapine *in vitro*. Using positron emission tomography (PET), we demonstrated that DCZ is rapidly brain penetrable, is apparently selective, and that doses for DREADD occupancy are 20- and 60-fold smaller than CNO and C21, respectively. Finally, we demonstrated that DCZ is capable of rapidly (<10 min post-injection) activating hM_3_Dq and hM_4_Di in both mice and monkeys without discernible off-target action. Thus, DCZ represents a potent and selective chemogenetic actuator for muscarinic-based DREADDs in mice and primates useful for a variety of *in vitro* and *in vivo* contexts with high translational potential.

## Results

### DCZ selectively binds DREADDs in vitro and in vivo

We first assessed the binding affinities of DCZ to hM_3_Dq and hM_4_Di, and compared them to those of clozapine, CNO and C21 using competition binding assays with [^3^H]quinuclidinyl benzilate (QNB) on HEK293T cells. DCZ had nM affinity for [^3^H]QNB-labelled hM_3_Dq and hM_4_Di (inhibition constants; ^hM3Dq^*K*_*i*_ = 6.3 nM; ^hM4Di^*K*_*i*_ = 4.2 nM), which was comparable to clozapine (^hM3Dq^*K*_*i*_ = 5.9 nM; ^hM4Di^*Ki* = 0.89). CNO and C21 were about 100- and 50-fold weaker, respectively (CNO, ^hM3Dq^*K*_*i*_ = 680 nM; ^hM4Di^*K*_*i*_ = 360 nM; C21, ^hM3Dq^*K*_*i*_ = 850 nM; ^hM4Di^*K*_*i*_ = 180 nM)(Table 1). Unlike clozapine (Fig. 1c and Supplementary Table 1), DCZ had negligible affinities for a large number of tested G protein-coupled receptors (GPCRs), ion channels or transporters (*K*_*i*_ s > 100 nM) and relatively low affinities for a few endogenous receptors including muscarinic acetylcholine (^hM1^*K*_*i*_ = 83 nM; ^hM5^*K*_*i*_ = 55 nM) and serotonin receptors (^5-HT2A^*K*_*i*_ = 87 nM)(Fig. 1b and Supplementary Table 1), representing at least an 8-fold antagonist binding selectivity for muscarinic DREADDs over its most potent endogenous targets.

We next examined whether DCZ enters the brain and binds to DREADDs *in vivo*. Here we performed PET with radiolabeled DCZ ([^11^C]DCZ) in two monkeys (see Supplementary Table 2 for a summary of the monkeys used in these experiments), which had received an adeno-associated virus type 2 vector (AAV2-CMV-hM_4_Di) injection, resulting in neuron-specific expression of hM_4_Di^6^ in the right putamen. An AAV2 control vector carrying the kappa-opioid-based DREADD (AAV2-CMV-KORD) was injected into the left putamen (Fig. 1e), as does not bind DCZ (not shown)^2^. Six weeks later, when stable expression is predicted^6^, the monkeys were scanned via PET after intravenous (i.v.) administration of a microdose of [^11^C]DCZ (Fig. 1f), revealing rapid brain penetrance of [^11^C]DCZ with high radioactive accumulation in the hM_4_Di-expressing putamen [standardized radio-ligand uptake value (SUV) at 85-90 min; 6.3 ± 0.4 for #209, 4.5 ± 0.8 for #212, mean ± sd]. Accumulation in the contralateral putamen was insignificant (2.4 ± 0.7 for #209, 1.7 ± 0.3 for #212)(Fig. 1g, for representative findings), in agreement with the high selectivity of DCZ for DREADDs *in vitro*. Low DCZ uptake was visible in surrounding cortical areas, potentially reflecting DCZ binding to endogenous receptors with low affinities. Using a single subject (#209), brain permeability and selectivity of DCZ were then directly compared to clozapine and CNO, two DREADD agonists whose radiolabeled forms were available; C21 radiolabeled with [^11^C] was unavailable. Consistent with our previous study^6^, PET imaging with microdoses of [^11^C]clozapine showed moderate uptake at the control putamen and cortical areas (Fig. 1h, left), likely reflecting its high binding affinity for many endogenous receptors (Fig. 1c and Supplementary Table 1), and increased uptake at the hM_4_Di-expressing putamen region presumably reflecting clozapine binding to hM_4_Di (Fig. 1h, left). PET imaging with [^11^C]CNO revealed a low brain signal, indicating modest brain permeability (Fig. 1i). Indeed, the whole brain concentration of [^11^C]CNO at 30 min was 0.14% of the injected dose, which was >40-fold lower than that of [^11^C]DCZ or [^11^C]clozapine (5.9% or 7.0%, respectively).

Dynamic regional radioactivity reflects radioligand localization in the vascular system and brain tissue in the initial phase, and ligand binding to tissue in the later phase. Here, the [^11^C]DCZ signal at the target region continued to increase, nearly reaching a plateau 90 min after radioligand injection, whereas the signal on the contralateral side increased moderately and then decayed rapidly (Fig. 1g, right). These regional differences in radioactivity were abolished by pretreatment with unlabeled DCZ (1 mg/kg, i.v.) (Supplementary Fig. 1a), and thus reflect displaceable ligand binding. We quantified specific [^11^C]DCZ binding to tissue using the cerebellum as reference region, where radioligand kinetics were unchanged by pretreatment (Supplementary Fig. 1a). On the parametric image representing specific binding, BP_ND_ (i.e., binding potential relative to non-displaceable uptake), high [^11^C]DCZ was localized to the hM_4_Di-vector injected side of the putamen, corresponding to the area where anti-hM_4_ post-mortem immunolabeling was found (Fig. 2a,b and Supplementary Fig. 2a), confirming that [^11^C]DCZ bound to hM_4_Di *in vivo*. In addition, high [^11^C]DCZ binding was also found in the projection target of the putamen — the substantia nigra — reflecting hM_4_Di expression at the axon terminals as confirmed by immunohistochemistry (Supplementary Fig. 2b).

**Fig. 2.**
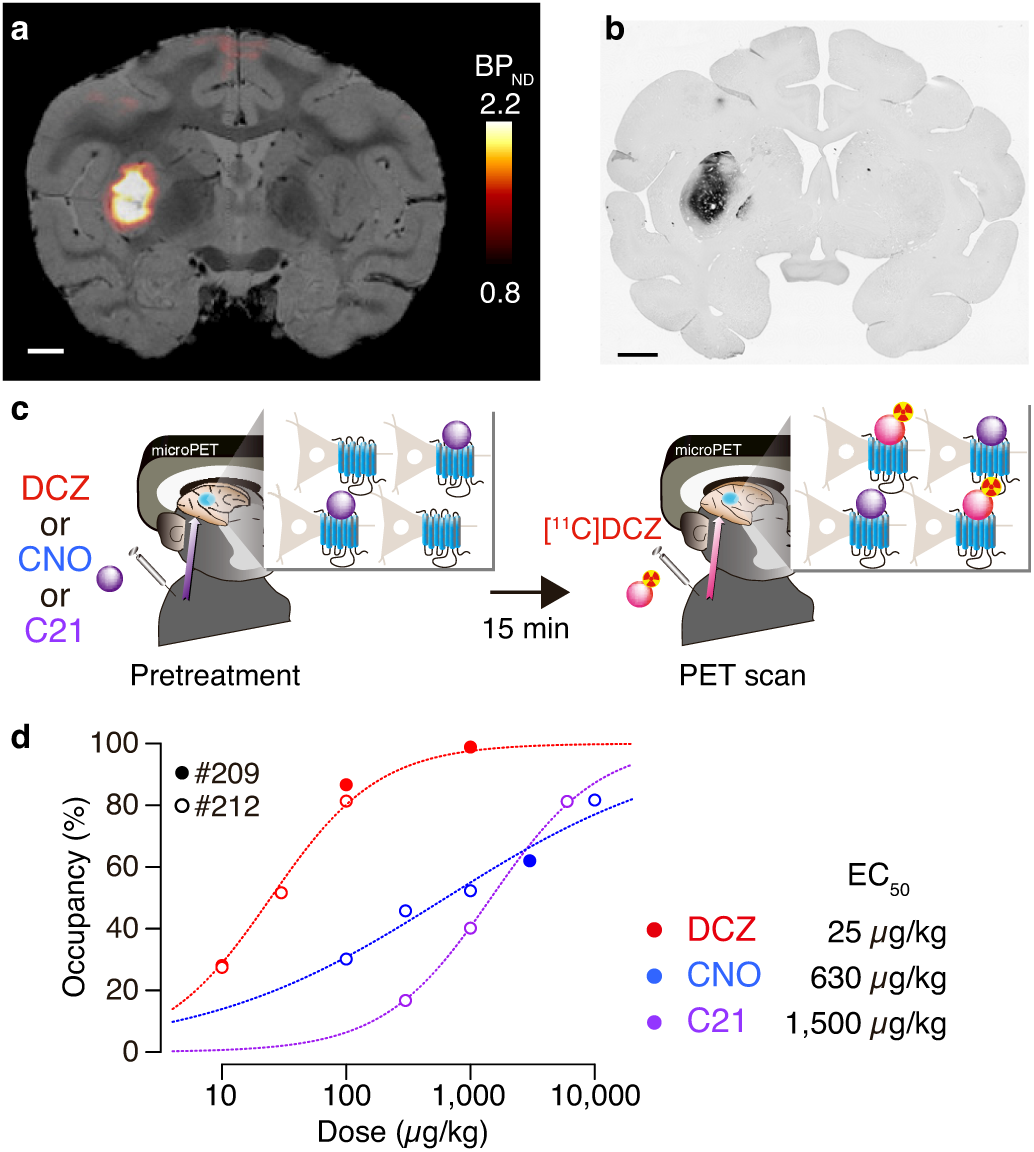
[^11^C]DCZ-PET visualizes DREADD expression and measures agonist dose–occupancy relationship. **a**, Coronal section of parametric [11C]DCZ-PET image of specific binding (BP_ND_) overlaying MR image of a monkey (#209) expressing hM_4_Di in the putamen. **b**, An anti-hM_4_ staining section corresponding to the image in **a**. Scale bars represent 5 mm. **c**, Illustration of occupancy study. Monkeys underwent [^11^C]DCZ-PET scan 15 min after i.v. injection of nonradiolabeled DCZ, CNO or C21. **d**, Occupancy of hM_4_Di plotted as a function of DCZ, CNO or C21 dose. Filled and open circles represent data obtained from monkeys #209 and #212, respectively. Dotted curves are the best-fit Hill equation for the data. ED_50_ indicates the agonist dose inducing 50% occupancy. Hill coefficient (n) and coefficient of determination (R^2^) as are follows: DCZ, n = 1, R^2^ = 0.99; CNO, n = 0.44, R^2^ = 0.96; C21, n = 1, R^2^ > 0.99.

To verify *in vivo* DCZ binding to DREADDs in mice, we performed [^11^C]DCZ-PET imaging with a transgenic mouse expressing hM_4_Di under the control of neuron-specific Thy-1 promoter^15^. [^11^C]DCZ uptake in the striatum and cortex was upregulated in transgenic mouse compared with wild-type mice (Supplementary Fig. 1b, left vs. center). In the parametric image, high [^11^C]DCZ binding was observed in the frontal and parietal cortices, hippocampus, and striatum, consistent with high hM_4_Di-expression as confirmed by immunofluorescence, whereas no discernible binding was seen in wild-type mouse brains (Supplementary Fig. 2c, d), implying low off-target binding at this dose. Taken together, our PET results suggest that DCZ rapidly enters the brain and selectively binds to hM_4_Di expressed in both mice and monkeys.

### Systemic low dose DCZ occupies hM4Di-DREADD in vivo

In previous chemogenetic studies, a range of systemic CNO doses was used to obtain behavioral effects, suggesting that the required dose may vary in a manner dependent on the target neuronal population, type of DREADD and level/localization of expression, as well as several other factors^16^. Similar to most drugs, at pre-saturation a higher agonist dose affords a stronger chemogenetic effect, but also has greater off-target potential. Although DCZ has lower levels of off-target binding than clozapine, finding the effective and safe dose range for systemic DCZ administration is critical for studies seeking selective DREADD manipulation. We previously used PET imaging to measure the relationship between CNO dose and the degree of hM_4_Di receptor occupancy, and provided a reasonable upper CNO dose (≤10 mg/kg); these doses successfully gave induced behavioral alterations in monkeys, while higher doses did not increase occupancy^6^. To quantify the relationship between DCZ dose and hM_4_Di occupancy, we performed [^11^C]DCZ-PET scans 15 min after i.v. bolus injection of non-radiolabeled DCZ (10, 30, 100 and 1,000 μg/kg) in two monkeys (Fig. 2c). With increasing doses of non-radiolabeled DCZ, specific binding of [^11^C]DCZ decreased at the hM_4_Di-vector injection site, while the decrease at the contralateral control site was limited. We determined the occupancy as a reduction of BP_ND_ at the target region over the control side relative to baseline. The relationship between the occupancy of hM_4_Di and DCZ dose was fit to a Hill equation with a coefficient equal to approximately 1 (Fig. 2d; Online Methods), indicating that DCZ binds non-cooperatively. The data also suggest that a reasonable dose range of DCZ for monkey studies is ≤100 μg/kg: occupancy increases linearly based on the logarithmic dose in this range, while higher doses would contribute to a minimal increase in occupancy as predicted based on standard receptor-ligand relationships.

Additional occupancy studies with CNO and C21 revealed that higher doses were required to occupy hM_4_Di (Fig. 2d). The dose-occupancy relationship of C21 indicated its binding property (Hill coefficient, n = 1) was similar to DCZ, while that of CNO (n = 0.44) might reflect complex kinetics of CNO metabolism and excretion. We estimated the dose required for 50% occupancy (ED_50_) for DCZ and compared it with those for CNO and C21: the ED_50_ for DCZ was 25 μg/kg, which was 24- and 60-fold smaller than for CNO (630 μg/kg) and C21 (1,500 μg/kg), respectively (Fig. 2d).

Pharmacokinetic (PK) studies demonstrated that DCZ administration (100 μg/kg, i.v.) yielded a maximum concentration of DCZ in CSF (∼10 nM) at 30 min post-injection which was maintained for at least 2 h in monkeys (Fig. 3a, right). A similar level of DCZ in CSF (∼10 nM) was observed at 60 min and later following intramuscular administration (Fig. 3b, right). The DCZ concentration was higher than its *K*_*i*_ value for hM_4_Di in antagonist radioligand binding studies (4.2 nM; Table 1) and far below its *K*_*i*_ value for numerous endogenous channels, receptors and transporters (> 50 nM; Fig. 1b). We did not detect any major metabolites of DCZ in CSF (Fig. 3a,b, right) in monkeys. Collectively, these results suggest that a low systemic dose of DCZ (i.e. 100 μg/kg) affords a sufficient concentration of DCZ for hM_4_Di-DREADD binding *in vivo* for at least 2 h without the detection of metabolites (Fig. 3d) in monkeys.

**Fig. 3.**
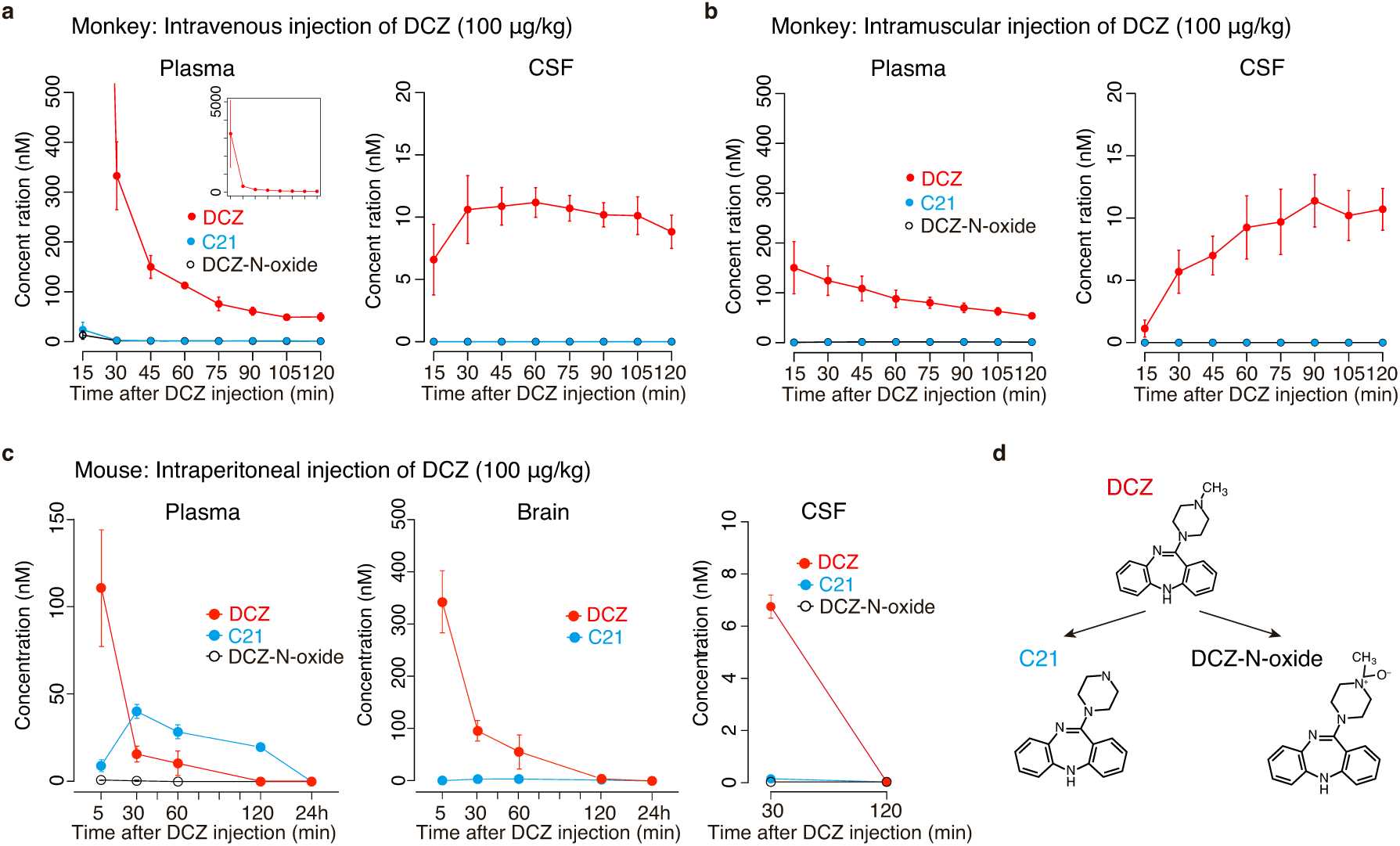
Time concentration profiles of DCZ and its metabolites in monkeys and mice. **a**, Plasma and CSF concentrations of DCZ and its major metabolites (C21 and DCZ-N-oxide) following its intravenous injection (100 μg/kg). Data were collected from 4 monkeys and are shown as mean ± sem. **b**, Same as a, but following intramuscular injection. **c**, Plasma, brain and CSF concentrations of DCZ, C21 and DCZ-N-oxide following intraperitoneal injection of DCZ (100 μg/kg). Data were collected from 3 or 4 mice and are shown as mean ± sem. **d**, Major routes of DCZ metabolism leading to C21 and DCZ-N-oxide and their structures.

We also found that DCZ was rapidly available in mouse brain following intraperitoneal (i.p.) administration (Fig. 3c). The DCZ concentration was ∼6-7 nM in CSF and ∼100 nM in brain tissue 30 min after administration (Fig. 3c). As the free (unbound) fraction of drug is key for determining the amount of drug that can bind to a target protein, we quantified brain protein binding. Such analysis revealed that the fraction of DCZ unbound to mouse brain tissue was about 7.5% (at 1 μM test concentration), indicating that a major fraction of DCZ bound nonspecifically to brain proteins as previously described for many CNS-active compounds^17,18^; the free DCZ concentration in CSF is comparable in mice (6-7 nM) and monkeys (∼10 nM). Unlike the PK profiles in monkeys, DCZ concentrations in mice declined rapidly and were undetectable at 2 h in either brain tissue or CSF. It is unsurprising that the PK properties of DCZ differ in mice and monkeys, as species differences in the PK properties of CNS active-drugs are well documented^19^. Although C21, DCZ’s desmethyl metabolite (Fig. 3d), was abundant in plasma, C21 was negligible in CSF (<0.2 nM, ∼ 3% of DCZ at 30 min, Fig. 3c).

### DCZ is a potent in vitro DREADD agonist

We next quantified the *in vitro* agonist activity for muscarinic-based DREADDs and compared its potency with several previously reported agonists. We first performed *in vitro* Bioluminescence Resonance Energy Transfer (BRET) assays, which directly measure agonist-induced G protein dissociation, the most proximal signaling event downstream of GPCR activation, with minimal potentiation due to so-called ‘receptor reserve’ or down-stream signal amplification^20^. Using BRET-based assays, DCZ was a potent hM_3_Dq agonist (EC_50_ = 0.13 nM; 95% confidence interval, 0.09-0.21) — comparable to clozapine [^hM3Dq^EC_50_ = 0.09 (0.06-0.15) nM], and about 40- and 100-fold more potent than C21 [^hM3Dq^EC_50_ = 5.2 (3.7-7.3) nM] and CNO [^hM3Dq^EC_50_ = 15 (8.8-25) nM], respectively (Fig. 4a). DCZ was also a potent agonist for hM_4_Di [^hM4Di^EC_50_ = 0.081 (0.042-0.16) nM], again comparable to clozapine [^hM4Di^EC_50_ = 0.051 (0.027-0.097) nM], but with about 30- and 90-fold greater potency than C21 [^hM4Di^EC_50_ = 2.6 (1.6-4.4) nM] and CNO [^hM4Di^EC_50_ = 7.3 (3.6-15) nM] (Fig. 4b), respectively.

**Fig. 4.**
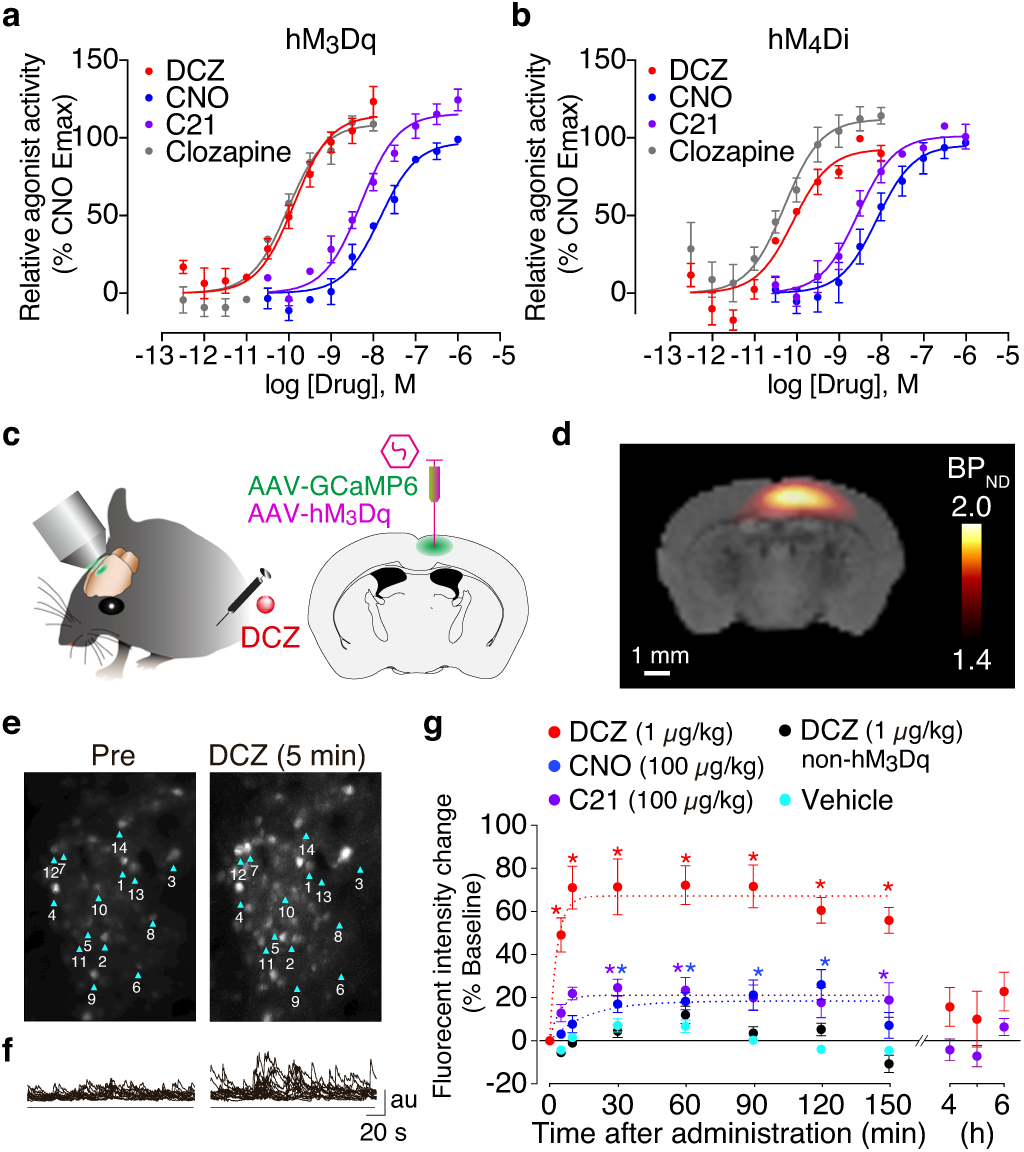
*In vitro* potency and *in vivo* efficacy of DCZ for DREADDs. **a**,**b**, BRET-based estimation of agonist potency and efficacy at hM_3_Dq and hM_4_Di on HEK293T cells. Shown are mean ± sem of n = 3 of independent biological replicates (each of which had duplicate technical replicates) of dose-response curves for hM_3_Dq using a Gq-based BRET sensor and hM_4_Di using a Gi1-based BRET sensor. **c**, Illustration of two-photon calcium imaging and location for co-injection of AAV2-CMV-hM_3_Dq and AAV-DJ-rSyn-GCaMP6s. **d**, Parametric image of specific binding of [11C]DCZ overlaying MR image. **e**, Time-averaged two-photon images of GCaMP6 of pre-injection (left) and 5 min post-DCZ injection (1 μg/kg, i.p.)(right). **f**, Raw fluorescence signals in 14 representative neurons from imaging in **e**. au: arbitrary units. **g**, Fluorescent intensity change (mean ± sem from baseline) as a function of post-injection time. Data were obtained from neurons (n = 58) in four hM_3_Dq-expressing mice and neurons (n = 30) in two non-hM_3_Dq mice. Curves represent exponential fits to the data. Asterisks represent significant difference (p < 0.01, one-way ANOVA with post-hoc Dunnett) from pre-injection.

Superior *in vitro* potency of DCZ over C21 and CNO was also demonstrated in a Ca^2+^ mobilization assay for hM_3_Dq^12^ and for inhibition of cAMP accumulation for hM_4_Di; importantly no effect of DCZ was observed in the absence of DREADD expression (Supplementary Fig. 3a-c). To further evaluate potential DCZ off-target agonist actions, we assessed its agonist activity at 318 endogenous GPCRs using an arrestin recruitment assay platform^21^. While DCZ was a potent agonist for hM_3_Dq and hM_4_Di, it did not display significant agonistic activity for any of the 318 tested wild-type GPCRs at <10 nM, the typical concentration that could be reached with the dose used (Supplementary Fig. 3d-l). Taken together, these results imply that DCZ’s agonism is apparently selective for hM_3_Dq and hM_4_Di.

### DCZ selectively and rapidly enhances neuronal activity via hM3Dq-DREADD in vivo

We next examined the DCZ’s agonist actions on DREADDs and its time course *in vivo* using two-photon calcium imaging in mice expressing hM_3_Dq. AAV vectors carrying the GCaMP6s genes (AAV-DJ-rSyn-GCaMP6s) and hM_3_Dq construct (AAV2-CMV-hM_3_Dq) were co-injected into the barrel cortex (Fig. 4c). At 28 days post-injection, *in vivo* hM_3_Dq expression was visualized by [^11^C]DCZ-PET as a high radioligand binding region at the injection site (Fig. 4d and Supplementary Fig. 1b, right). Given its high potency *in vitro*, we used DCZ at a dose of 1 μg/kg. Shortly after i.p. injections of DCZ, transient and repetitive increases in fluorescence signals were observed in the soma of hM_3_Dq-expressing neurons (Figs. 4e,f). On average, relative signal changes from baseline (ΔF/F) rapidly increased (τ = 3 min) and became significant at 5 min post-injection (p < 0.01, one-way ANOVA with post-hoc Dunnett test). ΔF/F achieved their peak at about 10 min, plateaued for at least 150 min, and then returned to the baseline levels at 4 h post-injection (Fig. 4g, red; p = 0.53). Vehicle injection did not change the activity of the same neuronal population (Fig. 4g, cyan). The DCZ-induced increases in fluorescent signals were apparently mediated by hM_3_Dq because DCZ did not alter the signal in mice without DREADD expression (Fig. 4g, black). Although DCZ induced strong activity changes, it did not cause long-term apparent desensitization to hM_3_Dq-positive neurons after recovery, since the neurons demonstrated a reproduced chemogenetic activation by the 2nd DCZ dose and a replicated response to whisker stimulation (Supplementary Fig. 4).

To compare the *in vivo* agonist efficacy of DCZ with those of other ligands, we used 100-fold higher doses (100 μg/kg, i.p.) of CNO and C21, because they require higher doses to occupy and activate DREADDs (cf. Figs. 2d and 4a). Although administration of CNO and C21 increased ΔF/F of hM_3_Dq-expressing neurons, their peak values were less than 40% of that of DCZ (Fig. 4g, blue and purple). Moreover, the kinetics of CNO were relatively slow (τ = 17 min), with the upward trend lasting for 30 min post-injection and then plateauing (Fig. 4g, blue), as observed in a previous electrophysiological study^22^. While C21 quickly increased ΔF/F (τ = 4 min), the increase became significant only at 30 min post-injection or later (Fig. 4g, purple). Thus, DCZ rapidly and reversibly activates hM_3_Dq-expressing neuronal populations in mice *in vivo* with higher efficacy and potency than C21 or CNO.

Rapid and reversible chemogenetic neuronal control was also observed in monkeys after systemic DCZ administration. A total of four monkeys were injected with AAV2-CMV-hM_3_Dq into the unilateral amygdala (Fig. 5a; one used for electrophysiology and the other three for activation imaging; see below). At 48-66 days post-injection, *in vivo* hM_3_Dq expression was visualized by [^11^C]DCZ-PET as a high radioligand binding region at the injection site (Fig. 5b), which was confirmed by immunohistochemistry (Supplementary Fig. 5). We recorded local field potential (LFP) from the hM_3_Dq-positive area of one monkey using a multi-site linear probe, the location of which was confirmed by CT-PET fusion image (Fig. 5b). To the best of our knowledge, there has been no prior report on a monkey chemogenetic study using hM_3_Dq. Therefore, we applied the same dose of DCZ (1 μg/kg) as in mice to avoid potential risks of excitotoxicity or seizure due to excessive neuronal activation. In the amygdala, LFP gamma band activity, which is known to correlate with firing of local populations of neurons^23^, increased significantly from baseline by administration of DCZ (t(26) = 7.3, p = 8.7 × 10^−8^, paired t-test), but not by vehicle (t(25) = 0.28, p = 0.78; Fig. 5c,d). As control, when we placed an electrode outside the hM_3_Dq-positive sites, significant changes in gamma power were not observed after DCZ administration (t(10) = 0.46, p = 0.65, Fig. 5d). LFP-gamma power rapidly increased (τ = 4 min) from baseline and became significant at 5 min after DCZ administration, and this significant increase lasted for at least 45 min (Fig. 5e, p < 0.01, one-way ANOVA with post-hoc Dunnett test).

**Fig. 5.**
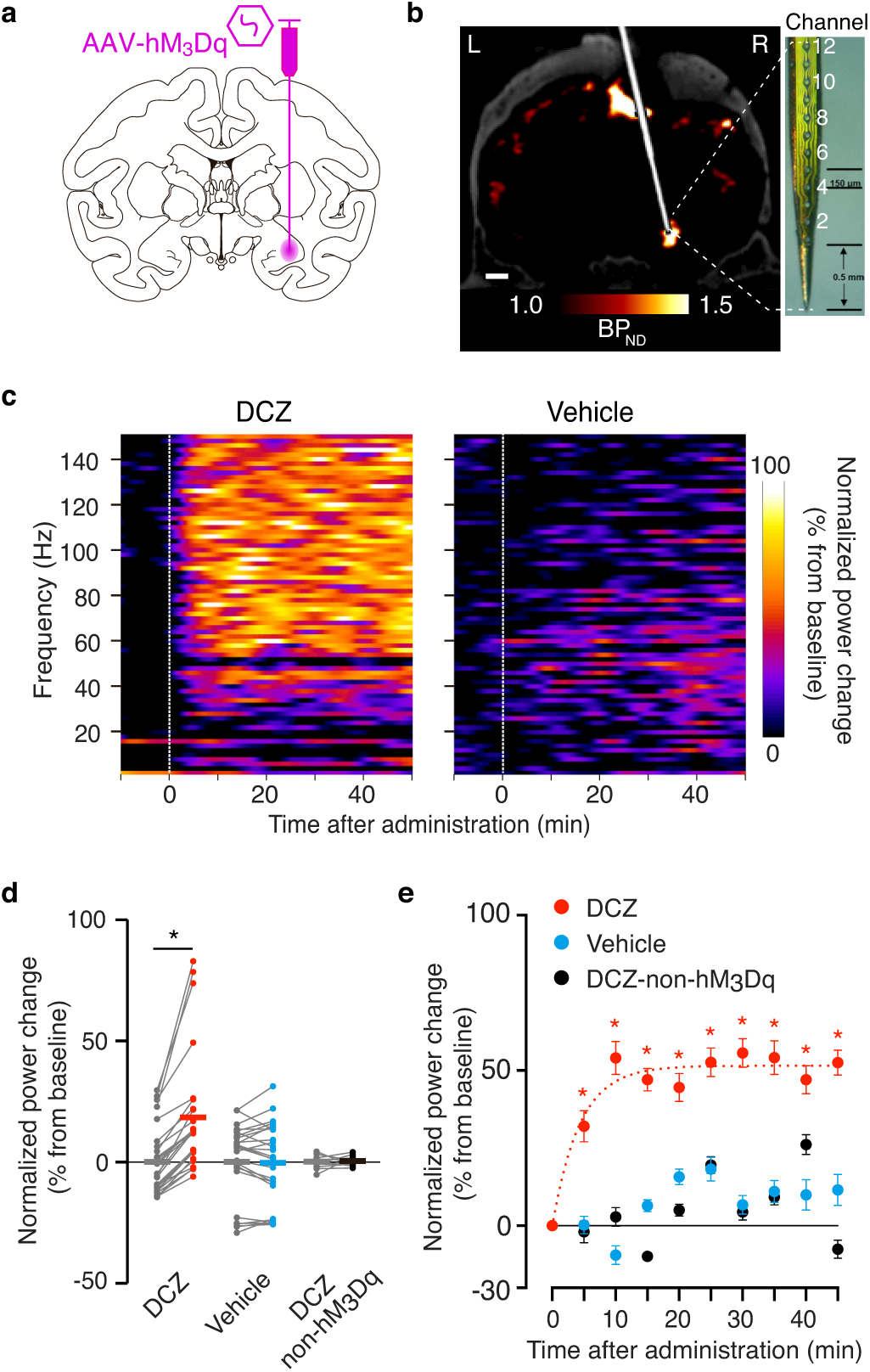
DCZ rapidly drives activation of hM_3_Dq-expressing neuronal population in monkey. **a**, Illustration representing viral vector (AAV2-CMV-hM_3_Dq) injection into one side of the amygdala. **b**, Coronal PET image overlaying CT image demonstrated that recording contacts on a multichannel electrode (shown on right) were located in an hM_3_Dq-expressing region, which was visualized as a high [^11^C]DCZ binding site. The high binding observed in the dorsomedial surface corresponds to a site of biological reaction within the recording chamber. Scale bar represents 5 mm. **c**, Representative LFP activity changes after DCZ (left) and vehicle administration (right). **d**, Normalized gamma (40 Hz) power change (mean ± sem from baseline) in 10-20 min after intravenous administration of reagents. Data were obtained from hM_3_Dq-positive region (DCZ and vehicle; n = 27 and 26 channels, respectively, acquired across 4 sessions) and non-hM_3_Dq expressing sites (DCZ-non-hM_3_Dq; n = 11 channels, acquired within 1 session). Gray bars indicate baseline. Asterisk represents significant difference from baseline. DCZ t(26) = 7.3, p = 8.7 × 10-8; vehicle t(25) = 0.28, p = 0.78; DCZ-non-hM_3_Dq t(10) = 0.46, p = 0.65. **e**, Plots represent normalized power change (mean ± sem from baseline) as a function of time from vehicle or DCZ administration. Curve represents exponential fit to the data. Asterisks represent significant difference (p < 0.01, one-way ANOVA with post-hoc Dunnett) from baseline. Data were obtained from one animal.

### DCZ selectively induces hM3Dq-mediated metabolic activity

We next performed a PET study with [^18^F]fluorodeoxyglucose (FDG) to examine whether DCZ induces dose-dependent and DREADD-selective changes in regional brain glucose metabolism, an index of brain neuronal/synaptic activation^24–26^. Three monkeys expressing hM_3_Dq in the unilateral amygdala underwent [^18^F]FDG PET imaging following systemic DCZ or vehicle injection (Fig. 6a-c). We found that FDG uptake at the hM_3_Dq-positive amygdala significantly increased after injections of 1- and 3-μg/kg DCZ doses (paired t-test, p < 0.01 for each dose), and this was reproducible without a reduction due to repetitive activations (Fig. 6d). The increase of FDG uptake occurred in a dose-dependent manner at the hM_3_Dq-positive amygdala region, while uptake at the contralateral control area was unchanged (one-way ANOVA, main effect of dose, F(2,14) = 25.06, p = 2.4 × 10^−5^; Fig. 6d). Voxel-wise statistical analysis further revealed that significant increase in FDG uptake after DCZ administration (3 μg/kg, i.v.) occurred exclusively at the hM_3_Dq-positive area (p < 0.001, uncorrected; Fig. 6b,c). In control monkeys without hM_3_Dq vector injection (non-DREADD; N = 2), administration of DCZ (1 or 100 μg/kg) did not change FDG uptake in the amygdala (one-way ANOVA, main effect of dose, F(2,12) = 2.29, p = 0.14; Fig. 6e). Voxel-wise statistical analysis demonstrated that no significant metabolic change was detected throughout the whole brain following a high dose of DCZ injection (100 μg/kg) in control monkeys, confirming that the off-target effect of DCZ is undetectable *in vivo* by this measure. These results suggest that DCZ induces a dose-dependent increase of chemogenetic neuronal excitation as measured by metabolic change with no significantly detectible off-target effects.

**Fig. 6.**
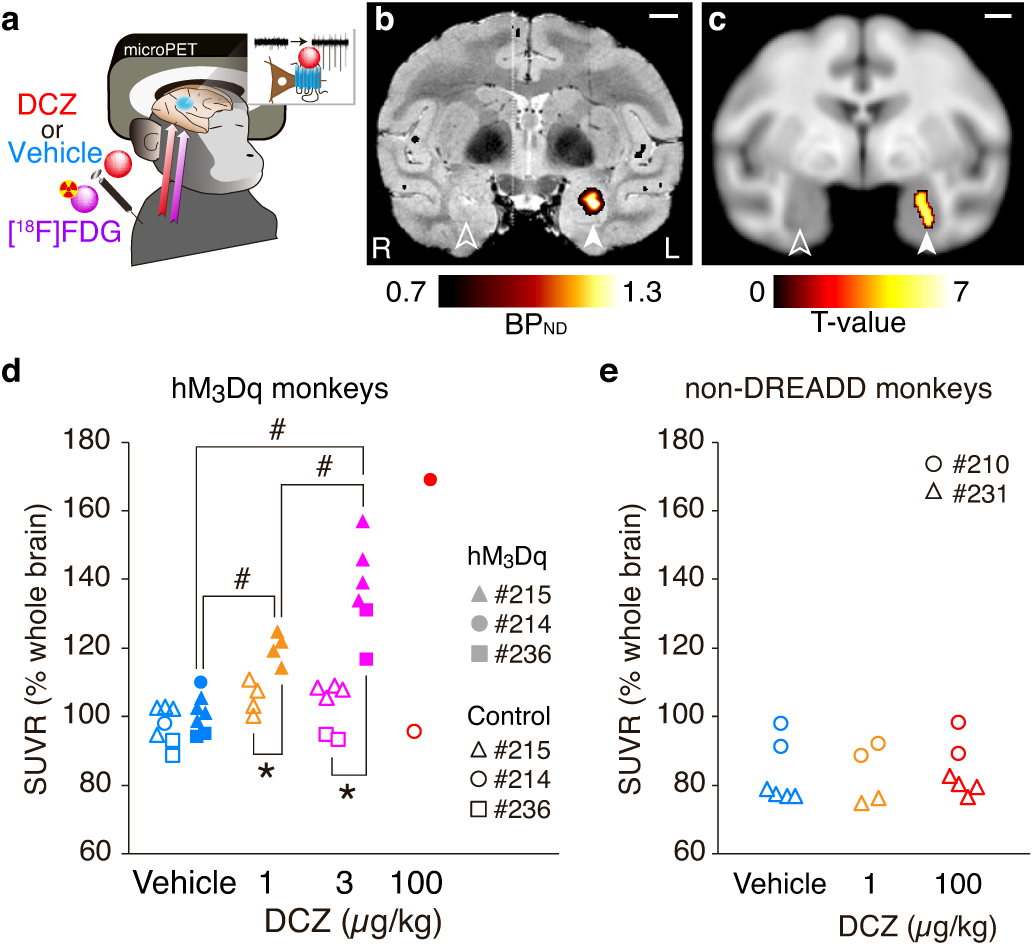
DCZ selectively induces metabolic changes in hM_3_Dq-expressing region in a dose-dependent manner. **a**, Illustration representing DREADD activation imaging with [^18^F]FDG. Following administration of DCZ at effective doses, [^18^F]FDG was injected and PET scan was performed to assess DREADD-induced brain metabolic change. **b**, Coronal PET image overlaying MR image representing specific [11C]DCZ binding (BP_ND_) in a monkey (#215). Filled and open arrowheads indicate the hM_3_Dq-expressing and control region in the amygdala, respectively. Scale bar represents 5 mm. **c**, Brain areas with significantly increased [^18^F]FDG uptake after DCZ injection (3 μg/kg, n=6 scans) compared with vehicle (n=6) overlay on MRI template (uncorrected p < 0.001, repeated measures one-way ANOVA, monkeys #215 and #236). Significance level is given as a t-value represented on a color scale. **d**, Relationship between the dose of DCZ administration and standardized FDG uptake (SUVR) in monkeys expressing hM_3_Dq in the amygdala. Filled and open symbols are the values at hM_3_Dq-expressing amygdala and contralateral control region, respectively. * p < 0.01, paired t-test (for 1 μg/kg DCZ, t(3) = 6.0, p = 0.009; for 3 μg/kg DCZ, t(5) = 9.7, p = 2.0 × 10^−4^). # p < 0.05, one-way ANOVA with post-hoc Tukey test (F(2,14) = 25.06, p = 2.4 × 10^−5^; vehicle vs. 1 μg/kg, p = 0.013; vehicle vs. 3 μg/kg, p = 0.03). **e**, Same as **d**, but for amygdala of control monkeys without hM_3_Dq-vector injection (non-DREADD monkeys; #210 and #231). There was no significant dose effect (one-way ANOVA, F(2,12) = 2.29, p = 0.14).

### DCZ selectively induces behavioral deficits in hM4Di-expressing monkeys

Finally, we sought to modify cognitive behavior using inhibitory DREADDs and DCZ. We targeted the prefrontal cortex (PFC), especially sectors around the principal sulcus corresponding to Brodmann’s area 46, which is responsible for spatial working memory and executive function. To implement chemogenetic silencing, two monkeys received multiple injections of an AAV-vector carrying hM_4_Di genes (AAV1-hSyn-hM_4_Di-IRES-AcGFP) bilaterally into the dorsal and ventral banks of the principal sulcus (Fig. 7a). [^11^C]DCZ-PET confirmed that *in vivo* hM_4_Di expression covered the bilateral target regions (Fig. 7b). We used a spatial delayed response task (Fig. 7c), which has been frequently employed as a sensitive probe of spatial working memory^7,27^. Compared with vehicle administration, intramuscular (i.m.) administration of DCZ (100 μg/kg) significantly impaired the performance of the delayed response task (two-way ANOVA with treatment × delay, main effect of treatment, F(1,24) = 63.53, p = 3.4 × 10^−8^, and F(1,24) = 188.5, p = 7.3 × 10^−13^ for monkeys #229 and #245, respectively; Fig. 7d). Impairment was more severe in the trials with longer delays (two-way ANOVA with treatment × delay, interaction, F(2,24) = 11.25, p = 3.6 × 10^−4^, and F(2,24) = 188.5, p = 4.3 × 10^−4^ for monkeys #229 and #245, respectively), suggesting that it was attributable to loss of working memory function. Impairment of the delayed response task appeared shortly after administration (<10 min) and lasted for at least 2 h (t-test, p < 0.05 for all test time periods; Fig. 7e), but disappeared at 24 h (two-way ANOVA with treatment × delay, main effect of treatment, vehicle vs 24 h after DCZ, F(1,24) = 0.60, p = 0.45, and F(1,24) = 0.94, p = 0.34 for monkeys #229 and #245, respectively; DCZ vs 24 h after DCZ, F(1,24) = 59.48, p = 6.0 × 10^−8^, and F(1,24) = 171.6, p = 2.0 × 10^−12^ for monkeys #229 and #245, respectively; Fig. 7d). The behavioral effect with DCZ was reproducible without attenuation due to repetitive DREADD activation (Fig. 7f). Without a screen during the delay period, DCZ did not affect the monkey’s performance, indicating that the impairment was unlikely attributable to deficits in motor function, visual perception or general motivation (Fig. 7g). We confirmed that DCZ alone did not produce any significant effects on spatial working memory in non-DREADD monkeys (N = 2, including #245, one of the two monkeys prior to vector injection; Fig. 7h). Additional behavioral examinations in three other non-DREADD monkeys using a reward-size task further corroborated that significant side-effect of DCZ injection (100 μg/kg, i.m.) was undetectable on motor or motivational functions (Supplementary Fig. 6). These results demonstrated that DCZ enables a rapidly and reversibly-induced behavioral change through activating muscarinic-based DREADDs without significant side effects.

**Fig. 7.**
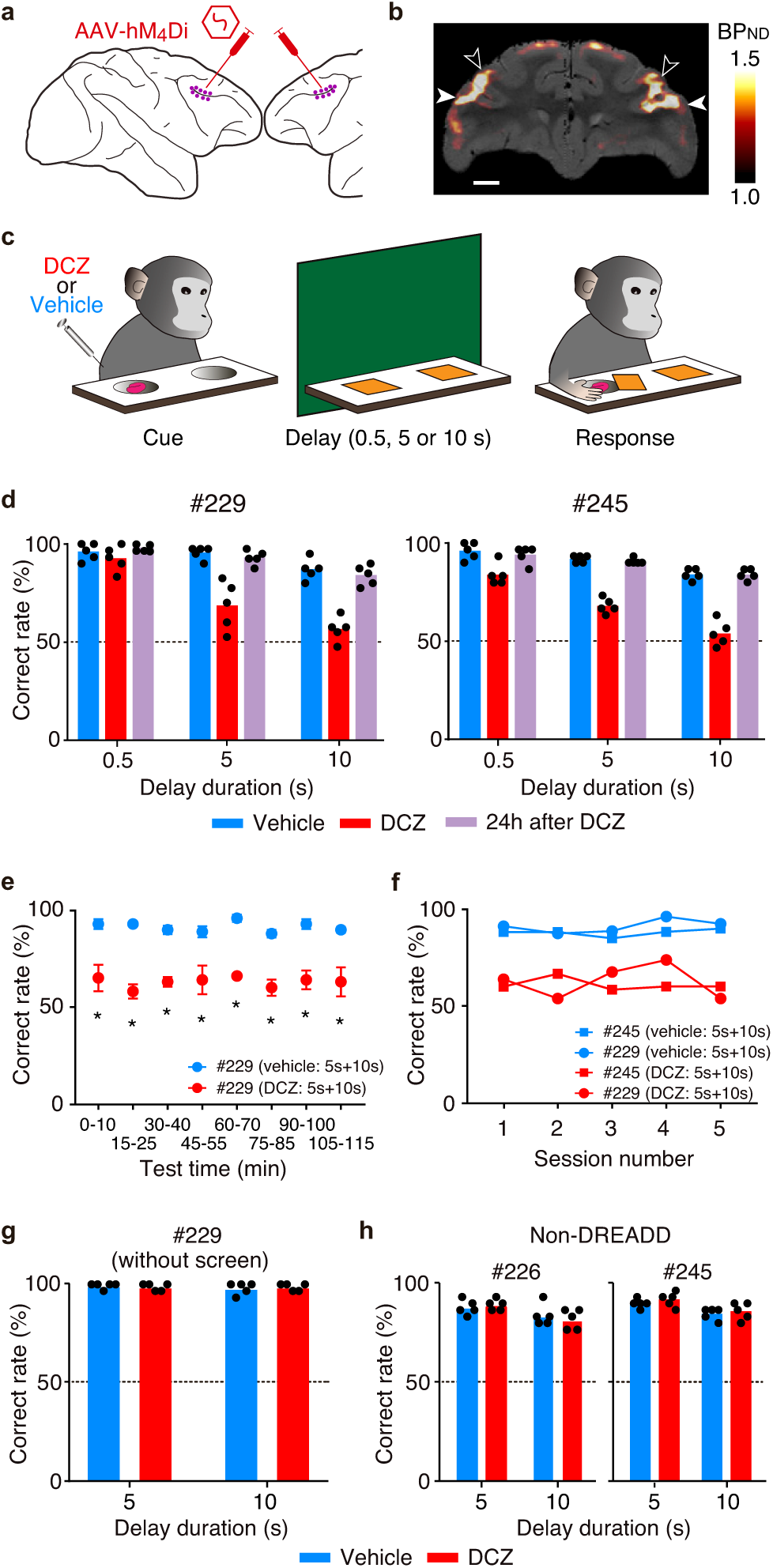
DCZ selectively and rapidly induces spatial working-memory deficit in monkeys expressing hM_4_Di in PFC. **a**, Illustration representing location of viral vector injections. AAV1-hSyn-hM_4_Di-IRES-AcGFP was injected bilaterally into the dorsal and ventral banks of the principal sulcus. **b**, Coronal parametric image of specific binding of [^11^C]DCZ overlaying MR image in a monkey (#229). Open and filled arrowheads represent the dorsal and ventral borders of the target regions, respectively. Scale bar represents 5 mm. **c**, Spatial delayed response task. After the monkey was shown the baited well (Cue), the screen was placed for 0.5, 5 or 10 s (Delay). The monkey obtained the food if they reached for the correct food-well and removed the cover plate (Response). **d**, Correct performance rates (mean ± sem) of spatial delayed response task with 0.5, 5 and 10 s delays after vehicle administration (n = 5 sessions; cyan), DCZ administration (100 μg/kg, i.m.; n = 5; red), and 24 h after DCZ administration (n = 5; purple) in two monkeys expressing hM_4_Di in PFC. Note that one of the monkeys (#245) was also examined prior to vector injection (see Supplementary Fig. 6c, right). The behavioral testing began immediately after an i.m. administration of either vehicle (2% DMSO in saline) or DCZ, except for 24 h after DCZ. Each dot represents the data in each daily session. **e**, Time course of DCZ effect. Correct performance rates of 5- and 10-s delays as a function of test time for vehicle and DCZ sessions (mean ± sem; n = 5) in monkey #229. The behavioral testings for 0-55 min and 60-115 min were conducted separately. Asterisks represent significant difference (t-test, p < 0.05). **f**, Correct performance rates (mean of 5- and 10-s delays) as a function of session number in two monkeys. **g**, Correct performance rates of the non-memory task after vehicle (n = 5 sessions; cyan) and DCZ administration (100 μg/kg, i.m.; n = 5; red) in monkey #229. No main effect of treatment, two-way ANOVA, F(1,16) = 1.1 × 10^−8^, p = 1.0. **h**, No significant effect of DCZ on delayed response task in two control monkeys without hM4Di-vector injection (two-way ANOVA, main effect of treatment, F(1,16) = 0.03, p = 0.87, and F(1,16) = 1.16, p = 0.30 for monkeys #226 and #245, respectively). Correct performance rates with 5- and 10-s delays after vehicle (n = 5 sessions; cyan) and DCZ administration (100 μg/kg, i.m.; n = 5; red) are shown. Note that the data of #245 were obtained prior to vector injection.

## Discussion

Here, we report that the new DREADD agonist DCZ represents an extremely potent, highly brain-penetrant, and selective actuator for hM_3_Dq- and hM_4_Di-DREADDs in both mice and monkeys. The properties of DCZ described here demonstrate that it can be adopted as a preferred, fast-acting chemogenetic actuator with minimal off-target actions, thereby enhancing opportunities for investigating the causal link between neuronal activity and behavior with high translational potential.

Several features of DCZ indicate its superiority as a chemogenetic actuator over prior agonists. First, DCZ exhibits DREADD-selective binding both *in vitro* and *in vivo*. Radioligand binding assays demonstrated that DCZ is selective for hM_3_Dq and hM_4_Di. PET imaging demonstrated that DCZ has substantially improved the selectivity compared with clozapine; DCZ has decreased off-target binding, while it retains similar potency at muscarinic DREADDs as clozapine. Second, DCZ exhibits good brain concentration profiles and biostability. PET imaging revealed that systemically administered DCZ occupied hM_4_Di-DREADDs with a similar occupancy to CNO or C21 at ∼20- or 60-fold smaller doses, respectively. Pharmacokinetic studies confirmed that DCZ is rapidly accumulated in mouse brains and monkey CSF, while its metabolites are negligible. Third, DCZ has high potency for muscarinic DREADDs. *In vitro* assays revealed that hM_3_Dq could be activated by substantially lower concentrations of DCZ, as compared to CNO and C21. DCZ did not display significant agonist potencies for any of the 318 off-target GPCRs tested; binding assays at a number of GPCRs, channels and transporters screened by the National Institute of Mental Health Psychoactive Drug Screening Program showed that DCZ has modest affinity (55 – 87 nM) for a few GPCRs, low affinity (100-1,000 nM) for several GPCRs and minimal activity (>1,000 nM) at most CNS targets tested. In short, DCZ has a higher selectivity for DREADDs than clozapine and is capable of occupying and activating DREADDs at a lower dose than CNO and C21, thus offering a greater effective window than the prior DREADD ligands.

CNO has repeatedly been shown to have relatively slow kinetics for neuronal activation via hM_3_Dq; following systemic administration, increases in activity began at around 5-10 min and reached a maximum at 45 min or later^22^. Because bath application of CNO immediately increased the firing rate of hM_3_Dq-expressing neurons in slice preparations^22^, the relatively slow activation appears to reflect the slow kinetics of CNO uptake in the brain. Moreover, the major effects on DREADDs may be exerted by its metabolite clozapine, rather than CNO itself^8^, where the metabolism is significant in rodents^8,9^ and in monkeys with slower delivery (e.g., subcutaneous injection^10^). Although the concentration of metabolized clozapine is minimal in monkeys with rapid intravenous delivery of CNO^6,28^, the brain uptake of CNO is considerably low (cf. Fig. 1i). In contrast, DCZ quickly penetrates into the brain and induces a rapid onset of excitability of hM_3_Dq-positive neuronal populations, as demonstrated by two-photon calcium imaging in mice and by electrophysiological recording in monkeys (cf. Figs. 4 and 5). Thus, DCZ combined with muscarinic-based DREADDs provides a superior platform for examining the effects of rapid and sustainable chemogenetic neuronal modulation. For example, single-unit or *in vivo* patch-clamp recording studies, where long-term (30-60 min) maintenance is technically demanding, would greatly benefit from the rapid action of the DCZ-DREADD system. Such a rapid chemogenetic action would also be valuable for future therapeutic applications, for instance, on-demand seizure attenuation^29^.

The present results indicate that DCZ can also be utilized for *in vivo* neuronal silencing by activating hM_4_Di, an inhibitory DREADD. Our data suggest that a reasonable upper limit of DCZ dose is 100 μg/kg (Fig. 2d), which allows a sufficient concentration of DCZ available for hM_4_Di-DREADD binding *in vivo* for at least 2 h in monkeys. Indeed, a 100 μg/kg dose of DCZ rapidly (<10 min) and reversibly induced spatial working memory deficits in monkeys expressing hM_4_Di in the PFC (Fig. 7). Given the high agonist potency of DCZ (Fig. 4b), smaller DCZ doses would induce similar or detectable behavioral deficits. With its high selectivity and potency, DCZ will provide potential orthogonality of the muscarinic DREADDs to other classes of chemogenetic tools^2,30^, providing opportunity for a multiplexed/bimodal control of physiological systems. Moreover, in combination with low-invasive targeted gene delivery methods such as ultrasonic BBB opening^31^, DCZ may open the potential for targeted, minimally invasive, and side-effect-free brain theranostics.

One of the technical difficulties in chemogenetic manipulations, especially for large animals, is the delivery of transgene specifically into a target neuronal population as well as the accomplishment of stable transgene expression throughout the experimental period. Previous studies have shown that PET imaging with [^11^C]clozapine or [^11^C]CNO can visualize hM_4_Di expression in living mice and monkeys, providing effective tools for non-invasive monitoring of gene expression^6,15^. However, because these ligands display poor DREADD selectivity or brain permeability, the application of DREADD-PET in monkeys was limited to visualization of hM_4_Di expression in the striatum. The present study has demonstrated that [^11^C]DCZ is an exceedingly useful PET ligand for visualizing both hM_3_Dq and hM_4_Di expression in cortical and subcortical areas of mice and monkeys, with improved specificity compared with [^11^C]clozapine (Fig. 1). The monitoring of transgene expression is beneficial for long-term behavioral studies and is of great advantage for conducting successful experiments (cf. Fig. 7). In addition to expression monitoring, PET imaging has allowed the elucidation of a relationship between a pharmacologically effective dose of DREADD agonists and DREADD occupancy *in vivo* (Fig. 2d). Moreover, it has also been demonstrated in our study that [^11^C]DCZ-PET enables us to visualize DREADDs expression at axon terminal sites (Supplementary Fig. 2), providing a powerful means for mapping projection areas for pathway-selective activity manipulation^32^, which can be induced by local infusion of DREADD agonist; DCZ may be suitable because of its high selectivity and efficacy.

Although PET imaging data have implied that DCZ has off-target binding in cortical areas possibly to endogenous receptors with low affinities (Fig. 1g and Supplementary Table 1), the agonist potency of DCZ at DREADDs is in the sub-nM range and, therefore, when used at a sufficient dose, occupancy on off-targets (∼50 nM or higher) can be predicted to be minimal. Indeed, our data further suggest that systemic administration of DCZ at 100 μg/kg or smaller doses would not induce discernible off-target effects on neuronal activity or behavior in monkeys not expressing the muscarinic-based DREADDs (cf. Fig. 6 and Supplementary Fig. 6). Since the variability of CSF concentration across subjects and the amount of metabolites for DCZ were much smaller than those for CNO (Fig. 3a)^6^, the potential baseline sensitivity to DCZ would be minimal. Therefore, DCZ provides a clear advantage over prior agonists due to increasing reliability by minimizing concerns about potential off-target effects. However, we cannot completely rule out unanticipated off-target effects because it is impossible to measure the activity of DCZ against every potential endogenous target. Therefore, as with all other neuromodulation technologies, control experiments—here using non-DREADD animals—are recommended. In mice, control experiments are essential since the current study did not examine an off-target action of DCZ at a high dose where C21, a DCZ metabolite, was detected at very low concentrations in CSF (<0.2 nM; Fig. 3c). Future studies will be required to determine appropriate DCZ doses and the effective time window for mouse chemogenetic experiments.

In conclusion, the characteristics of DCZ described here—its high selectivity, high brain-penetrability, and biostability—facilitate the rapid and selective modulation of neuronal activity and behavior with muscarinic-based DREADDs in living animals. Given the potential drawbacks of prior DREADD agonists, DCZ will provide clear benefits for many users of muscarinic DREADDs, with increasing reliability by removing concerns about potential off-target responses.

## Acknowledgements

We thank R. Suma, J. Kamei, R. Yamaguchi, Y. Matsuda, Y. Sugii, A. Maruyama, T. Okauchi, T. Kokufuta, Y. Iwasawa, T. Watanabe, A. Tanizawa, S. Shibata, N. Nitta, Y. Ozawa, M. Fujiwara, M. Nakano, T. Minamihisamatsu, S. Uchida and S. Sasaki for their technical assistance. We also thank S. Hiura for 3D printing of grids. This study was supported by MEXT/JSPS KAKENHI Grant Numbers JP15H05917, JP15K12772 and JP18H04037 (to TM), JP16H02454 (to MTakada), JP19K08138 (to YN), and JP18H05018, JP19K07811 and JP20H04596 (to NM), by AMED Grant Numbers JP19dm0107146 (to TM), JP19dm0207003 (to MTakada), JP19dm0107094 and JP18dm0207007 (to TS), JP19dm0307021(to KI), JP19dm0307007 (to TH), and JP20dm0207072 (to MH), by JST PRESTO Grant Number JPMJPR1683 (to KI), by QST President’s Strategic Grant (Creative Research)(to NM), by the Cooperative Research Program at PRI, Kyoto Univ, by National Bio-Resource Project ‘‘Japanese Monkeys’’ of MEXT, Japan, and by U24DK116195, the NIMH Psychoactive Drug Screening Program and the Michael Hooker Distinguished Professorship to BLR.

## Author Contributions

Conceptualization, T.M.; Formal Analysis, Y.N., N.M., K.O., K.M., and T.M.; Investigation, Y.N., N.M., H.T., Y.H., K.O., B.J., M.Takahashi, X.-P.H., S.T.S., J.F.D., X.Y., T.U., J.G.E., J.L., K.K., C.S., M.O., and M.S.; Resources, B.J., X.Y., J.L., K.I., M.Takada, and J.J.; Writing – original draft, Y.N., N.M., and T.M.; Visualization, Y.N., N.M., H.T., Y.H., K.O., B.J., K.M. and T.M.; Supervision, M.-R.Z., T.S., M.Takada, M.H., J.J., B.L.R., and T.M.; Project Administration, B.L.R., and T.M.; Funding Acquisition, Y.N., N.M., T.H., K.I., M.Takada, T.S., M.H., J.J., B.L.R., and T.M.; Writing – review & editing, all authors.

## Methods

### Subjects

All experimental procedures involving animals were carried out in accordance with the Guide for the Care and Use of Laboratory Animals (National Research Council of the US National Academy of Sciences) and were approved by the Animal Ethics Committee of the National Institutes for Quantum and Radiological Science and Technology. A total of 16 macaque monkeys [7 Rhesus (*Macaca mulatta*), and 9 Japanese monkeys (*Macaca fuscata*); 11 males, 5 females; 2.8-8.0 kg; age 4-10 years] were used (a summary of subjects used in the experiments described in Supplementary Table 2). The monkeys were kept in individual primate cages in an air-conditioned room. A standard diet, supplementary fruits/vegetables and a tablet of vitamin C (200 mg) were provided daily. Adult hM4Di transgenic mice with C57BL/6j background^15^ (male, age >12 weeks) and age-matched non-Tg littermates or wild-type C57BL/6j mice (male, age >12 weeks, Japan SLC Inc., Hamamatsu, Japan) were used. All mice were maintained in a 12-h light/dark cycle with ad libitum access to standard diet and water.

### Viral vector production

AAV2 (AAV2-CMV-hM_4_Di, AAV2-CMV-KORD, AAV2-CMV-hM_3_Dq, AAV2-CMV-AcGFP; Figs. 1d, 4c, 5a) and AAV1 (AAV1-hSyn-hM_4_Di-IRES-AcGFP; Fig. 7a) vectors were produced by helper-free triple transfection procedure, which was purified by affinity chromatography (GE Healthcare, Chicago, USA). Viral titer was determined by quantitative PCR using Taq-Man technology (Life Technologies, Waltham, USA). Transfer plasmid was constructed by inserting cDNA fragment and WPRE sequence into an AAV backbone plasmid (pAAV-CMV, Stratagene, San Diego, USA). For production of an AAV-DJ vector (AAV-DJ-rSyn-GCaMP6s), a transfer plasmid containing rat Synapsin promoter and cDNA encoding GCaMP6s (Addgene plasmid #40753) was assembled and transfected with helper-free DJ plasmids (Cell Biolabs, San Diego, USA). Viral particles were purified by HiTrap heparin column (GE Healthcare). Viral titer was determined by AAVpro® Titration kit ver2 (TaKaRa).

### Surgical procedures and viral vector injections

In monkeys, surgeries were performed under aseptic conditions in a fully equipped operating suite. We monitored body temperature, heart rate, SpO_2_ and tidal CO_2_ throughout all surgical procedures. Monkeys were immobilized by intramuscular (i.m.) injection of ketamine (5-10 mg/kg) and xylazine (0.2-0.5 mg/kg) and intubated with an endotracheal tube. Anesthesia was maintained with isoflurane (1-3%, to effect). Prior to surgery in monkeys, magnetic resonance (MR) imaging (7 tesla 400mm/SS system, NIRS/KOBELCO/Brucker) and X-ray computed tomography (CT) scans (Accuitomo170, J. MORITA CO., Kyoto, Japan) were performed under anesthesia (continuous infusion of propofol 0.2-0.6 mg/kg/min, i.v.). Overlay MR and CT images were created using PMOD® image analysis software (PMOD Technologies Ltd, Zurich, Switzerland) to estimate stereotaxic coordinates of target brain structures.

For subcortical injections, monkeys underwent a surgical procedure to open burr holes (∼8 mm diameter) for the injection needle. Viruses were pressure-injected by 10-μL Hamilton syringe (Model 1701 RN, Hamilton Company, Reno, USA) with a 30-gauge injection needle and a fused silica capillary (450 μm OD) to create a ‘step’ about 500 μm away from the needle tip to minimize backflow. The Hamilton syringe was mounted into a motorized microinjector (Legato130, KD Scientific or UMP3T-2, WPI, Sarasota, USA) that was held by manipulator (Model 1460, David Kopf, Ltd., Tujunga, USA) on the stereotaxic frame. After the dura mater was opened about 3 mm, the injection needle was inserted into the brain and slowly moved down 2 mm beyond the target and then kept stationary for 5 min, after which it was pulled up to the target location. The injection speed was set at 0.25-0.5 μL/min. After each injection, the needle remained *in situ* for 15 min to minimize backflow along the needle. Two monkeys (#209 and #212) had co-injections of AAV vectors (total 6 μL; 3 μL × 2 different depths in a track) carrying hM_4_Di construct (AAV2-CMV-hM_4_Di, #209: 1.8 × 10e^13^ and #212: 2.6 × 10e^13^ particles/mL) and AcGFP genes (AAV2-CMV-AcGFP, 0.7 × 10e^13^ particles/mL) in one side of the putamen and co-injections of AAV vectors (total 6 μL; 3 μL × 2 different depths in a track) carrying kappa-opioid based DREADD construct (AAV2-CMV-KORD; 1.8 × 10e^12^ particles/mL) and AcGFP (AAV2-CMV-AcGFP, 0.7 × 10e^13^ particles/mL) in the other side. Four monkeys (#214, #215, #233, #236) had co-injections of AAV vectors (total 6 μL; 3 μL × 2 tracks, 2 mm apart rostrocaudally) carrying hM_3_Dq construct (AAV2-CMV-hM_3_Dq; 1.2 × 10e^13^ particles/mL) and AcGFP genes (AAV2-CMV-AcGFP, 0.7 × 10e^13^ particles/mL) into the left (#214 and #215) or right (#233, #236) amygdala.

Two monkeys (#229, #245) had AAV1-hSyn-hM_4_Di-IRES-AcGFP (3.8 × 10e^13^ particles/mL) injected into the bilateral prefrontal cortex (Brodmann’s area 46). After retracting the skin and galea, the frontal cortex was exposed by removing a bone flap and reflecting the dura mater. Handheld injections were made under visual guidance through an operating microscope (Leica M220, Leica Microsystems GmbH, Wetzlar, Germany), with care taken to place the beveled tip of the Hamilton syringe containing the viral vector at an oblique angle to the brain surface. One person inserted the needle into the intended area of injection and another person pressed the plunger to expel approximately 1 μL. Nine tracks were injected in each hemisphere; one was located 1 mm posterior to the caudal tip of the principal sulcus, and the others were located along the dorsal (4 tracks) and ventral (4 tracks) bank of the principal sulcus posterior to the rostral tip of the ascending limb of the arcuate sulcus (Fig. 7a). Viral vectors were injected at 3 to 5 μL per track depending on the depth. Total amounts of viral aliquots injected into the right and left hemispheres were 35 and 37 μL for #229, and 44 and 40 μL for #245, respectively.

In mice, the animals were anesthetized with a mixture of air, oxygen, and isoflurane (3– 5% for induction and 2% for surgery) via a facemask, and a cranial window (3-4 mm in diameter) was placed over the left somatosensory cortex, centered at 1.8 mm caudal and 2.5 mm lateral from the bregma, according to the ‘Seylaz-Tomita method’^33^. On the day of cranial window surgery, AAV vectors carrying GCaMP6 genes (AAV-DJ-rSyn-GCaMP6, 3.5 × 10e^11^ particles/mL) and hM_3_Dq construct (AAV2-CMV-hM_3_Dq, 1.5 × 10e^13^ particles/mL) were co-injected into the barrel cortex using glass needles. A custom metal plate was affixed to the skull with a 7-mm-diameter hole centered over the cranial window. The method for preparing the chronic cranial window was previously reported in detail^34^.

### Radioligand competition binding assays

Radioligand binding assays with membrane preparations to determine binding affinity were carried out by the National Institute of Mental Health’s Psychoactive Drug Screening Program (NIMH PDSP) (https://pdsp.unc.edu/). Detailed assay protocols are available at the NIMH PDSP website (http://pdspdb.unc.edu/pdspWeb/?site=assays). NIMH PDSP is directed by Bryan L Roth, MD, Ph.D., University of North Carolina at Chapel Hill, North Carolina, and Program Officer Jamie Driscoll at NIMH, Bethesda, USA.

### Bioluminescence Resonance Energy (BRET) experiments

HEK293T cells split into 10-cm plates and maintained in Dulbecco’s Modified Eagle Medium (DMEM) supplemented with 10% fetal bovine serum (FBS) and 1% penicillin-streptomycin (pen-strep) were co-transfected with 1 μg each of hM_3_D or hM_4_D and Gαq-RLuc8 or Gαi1-RLuc8, respectively, and 1 μg each of Gβ1 and Gγ2-GFP2 using TransIT-2020 (Mirus) as transfection reagent. For negative controls, 1 μg of pcDNA was transfected in place of hM_3_D or hM_4_D. After at least 12 h, cells were plated in poly-D-lysine-coated white 96-well microplates (Greiner) in 100 μL of DMEM supplemented with 1% dialyzed FBS and 1% pen-strep at a density of 50,000 cells per well. After at least additional 12 h, media were aspirated and replaced with 60 μL of assay buffer (1x Hanks’ Balance Salt Solution, 20 mM HEPES, pH 7.40). Next, 10 μL of 50 μM coelenterazine 400a (NanoLight Technology) was added. After a 5-min incubation, 30 μL of 3x drug (in assay buffer containing 0.3 mg/mL ascorbic acid and 0.3% bovine serum album) was added. After an additional 5-min incubation, plates were read for luminescence with a Mithras LB 940 multimodal microplate reader (Berthold Technologies). Data were analyzed by GraphPad Prism 7.0 using the built-in dose-response function, and were normalized to the responses produced by CNO. In the presence of RLuc8 substrate coelenterazine, proximal and properly oriented alpha-RLuc8 and gamma-GFP2 produce BRET that reflects the equilibrium association of the heterotrimeric G protein under a given set of conditions. Dissociation of the heterotrimer consequent to receptor activation decreases BRET, and the extent of this decrease is dose-dependent. Since this assay directly measures G protein dissociation, which is the most proximal signaling event downstream of GPCR activation, measurements made by this assay are relatively insensitive to receptor reserve and signal amplification (see Olsen et al^20^, for details regarding BRET assays and construct design).

### In vitro cAMP assays

G_i_-mediated inhibition of cAMP production assays were performed in transiently transfected HEK293T cells. Briefly, HEK293T cells, transfected overnight with GloSensor plasmid (Promega) and receptor DNAs, were plated (10-15,000 cells/40 μL per well) in poly-L-lysine-coated white 384-well clear-bottom cell culture plates in DMEM with 1% dialyzed FBS. After 16-20 h, cells were removed from medium and stimulated with ligands prepared in HBSS (Hank’s balanced salt solution), 20 mM HEPES, 0.1% BSA, pH 7.4 for 15 min, followed by 0.1 μM isoproterenol (final concentration) in GloSensor reagent. Luminescence was read on a Wallac TriLux Microbeta counter (PerkinElmer). Results were normalized to the CNO activity and analyzed using the built-in dose-response function in GraphPad Prism 7.0.

### In vitro Ca2+ mobilization (FLIPR) assays

HEK293T cells stably expressing hM_3_Dq or hM_3_ receptors were used for Gq-mediated calcium mobilization assays. Assays were performed according to published procedures^12^. More detailed assay protocols are available at the NIMH PDSP website (http://pdspdb.unc.edu/pdspWeb/?site=assays).

### GPCRome screening (PRESTO-Tango) assays

Potential agonist activity at human GPCRome was measured using the PRESTO-Tango assay as published^21^.

### Brain protein binding assays

The unbound fraction of DCZ in brain tissue was measured by an equilibrium dialysis method. PBS solutions containing mouse brain homogenates (20%) and DCZ (final concentrations of 10, 100 or 1,000 nM) were added to 96-well plates (Equilibrium Dialyzer MW10K; HARVARD APPARATUS), followed by equilibrium dialysis (22 h at 37°C). A certain amount of methanol solution containing sulfaphenazole was added as an internal standard to the separated filtrate, brain homogenate fractions and pre-dialyzed samples, and centrifuged (1,700g, 10 min). The supernatant fractions were applied to LC-MS/MS to measure DCZ concentration. The average fraction of unbound brain tissue was calculated for each concentration as reported in the previous publication^35^.

### Drug administration

DCZ (HY-42110, MedChemExpress or synthesized in house, see below) was dissolved in 1-2% of dimethyl sulfoxide (DMSO) in saline to a final volume of 0.1 mL/kg. CNO (Toronto Research, North York, Canada) was dissolved in 2.5% of DMSO in saline to a final volume of 0.1 mL/kg. C21 (Compound 21, TOCRIS, Bristol, UK) was dissolved in 2% of DMSO in distilled water to a final volume of 0.1 mL/kg. For plasma/CSF analysis, a 23-gauge catheter was placed in the saphenous vein or the spinal canal for acute sampling while the monkey was under ketamine and xylazine anesthesia. DCZ (100 μg/kg) was administered at a rate of 0.2 mL/s intravenously via catheter. For PET blocking and occupancy studies, DCZ (10, 30, 100 or 1,000 μg/kg), CNO solution (100, 300, 1,000, 3,000 or 10,000 μg/kg) or C21 (300, 1,000 or 6,000 μg/kg) was administered intravenously via a saphenous vein catheter 1-15 min before PET imaging. For FDG-PET study, DCZ (1, 3 or 100 μg/kg) solution or vehicle was administered intravenously 1 min before PET imaging. Fresh solutions were prepared on the day of usage.

### Synthesis of DCZ

1-methylpiperazine (0.33 mL, 2.97 mmol) was added to a solution of 11-chloro-5*H*-dibenzo[*b,e*][1,4]diazepine (0.22 g, 0.96 mmol)^12^ in toluene (5 mL). The resulting solution was refluxed for 2 h. After cooling down to room temperature, the reaction was concentrated. The resulting residue was purified by silica gel flash column chromatography with 0−10% MeOH in CH_2_Cl_2_ to give the desired product (0.20 g, yield 70%). ^1^H NMR (800 MHz, CD3OD) δ7.69 – 7.64 (m, 2H), 7.41 (d, J = 7.9 Hz, 1H), 7.33 – 7.29 (m, 2H), 7.27 (t, *J* = 7.6 Hz, 1H), 7.19 – 7.13 (m, 2H), 4.40 – 3.80 (br, 4H), 3.80 – 3.50 (br, 4H), 3.06 (s, 3H). HRMS calcd. for C_18_H_21_N_4_: 293.1761; found: 293.1720 [M + H]^+^.

### Radiosynthesis

[^11^C]DCZ was produced as follows. An automated multi-purpose synthesizer developed in house was used for the present radiosynthesis. The cyclotron-produced [^11^C]CO_2_ was converted to [^11^C]CH_3_I, which was distilled and sent through a silver trifluoromethanesulfonate glass tube under N_2_ gas flow to yield [^11^C]methyl trifluoromethanesulfonate ([^11^C]CH_3_OTf). [^11^C]CH_3_OTf was then introduced to a reaction vial containing desmethyl precursor (C21, 0.2 mg) in dichloromethane (0.3 mL) under room temperature. The reaction mixture was kept at room temperature for 5 min. HPLC purification was completed on an X-Bridge C_18_ column (10 mm i.d. × 250 mm, Waters, Milford, USA) using CH_3_CN/H_2_O/Et_3_N (40/60/0.1%) at 5.0 mL/min. The radioactive fraction corresponding to [^11^C]DCZ (t_R_: 9.5 min) was collected and formulated to obtain an injectable solution. Synthetic time was about 40 min from the end of bombardment with an averaged radiochemical yield (decay-corrected) of 76.2% based on [^11^C]CO_2_ (n = 16). Radiochemical purity was assayed by analytical HPLC (column: CAPCELL PAK C18 UG120 S5, 4.6mmID, 150mm length, Shiseido, Tokyo, Japan) UV at 254 nm; mobile phase: CH_3_CN/H_2_O = 40/60 (Et_3_N 0.1%). Radiochemical purity and molar activity of [^11^C]DCZ were >98% and 130 ± 45 GBq /μmol (n = 16), respectively.

[^11^C]clozapine was radiosynthesized from desmethylclozapine by ^11^C-methylation using ^11^C-methyl triflate based on the previously described protocol^36^, and its radiochemical purity and specific radioactivity at the end of synthesis exceeded 95% and 37 GBq/μmol, respectively. [^18^F]FDG was purchased from Nihon Medi-Physics Co., Ltd. (Tokyo, Japan).

### PET imaging

PET scans were performed using microPET Focus 220 scanner (Siemens Medical Solutions USA, Malvern, USA). Anesthesia was performed as follows: a mouse was anesthetized with 1-3% isoflurane and a monkey was immobilized by intramuscular injection of ketamine (5-10 mg/kg) and xylazine (0.2-0.5 mg/kg) and then maintained in anesthetized condition with isoflurane (1-3%) during all PET procedures. Transmission scans were performed for about 20 min with a Ge-68 source for monkey scans. Emission scans were acquired in 3D list mode with an energy window of 350–750 keV after intravenous bolus injection of [^11^C]DCZ (31.8-46.4 MBq for mice and 276.9-386.9 MBq for monkeys), [^11^C]clozapine (287.4-353.8 MBq), [^11^C]CNO (354.6 MBq) or [^18^F]FDG (157.6-358.5 MBq). The actual injected dose depended on the specific radioactivity and the weight of the subjects (for example, <0.8 μg/kg for monkey #209). Emission data acquisition lasted 90 min for [^11^C]DCZ, [^11^C]clozapine and [^11^C]CNO scans and 120 min for [^18^F]FDG scans. All list-mode data were sorted into three-dimensional sinograms, which were then Fourier-rebinned into two-dimensional sinograms (frames × minutes: 5 × 1, 5 × 2, 5 × 3, and 12 × 5 for [^11^C]DCZ and [^11^C]clozapine, 24 × 5 for [^18^F]FDG). Images were thereafter reconstructed with filtered back-projection using a Hanning filter cut-off at a Nyquist frequency (0.5 mm^−1^). Standardized uptake value (SUV) was calculated using PMOD® image analysis software as the regional concentration of radioactivity averaged across the specific time window after injection of the radioligand. Volumes of interest (VOIs) were placed using PMOD® image analysis software with reference to the MR image of individual monkeys or the mouse brain template generated as described previously^15^. To obtain accurate registrations between PET and MR images, the rigid matching function of PMOD® (PFUS) was used. For this PET-MR registration, maximum errors in translation and rotation of human brain data were 1.67 mm and 0.6 degrees, respectively^37^, despite the fact that no data were available for monkey brains. In addition to the registration error, there should be spatial limitation of PET imaging of the “positive region” due to the signal spread from the source. Based on our *in vivo* and *in vitro* cross reference, we approximated that the limitation was <2 mm. In FDG studies with hM_3_Dq-expressing monkeys, VOI for the hM_3_Dq-positive region was defined as the area where the BP_ND_ value of [^11^C]DCZ was higher than 1.0, while that of the control region was placed at the corresponding contralateral side.

### Two-photon laser-scanning microscopy

Awake mice were placed on a custom-made apparatus, and real-time imaging was conducted by two-photon microscopy (TCS-SP5 MP, Leica Microsystems GmbH) with an excitation wavelength of 900 nm. An emission signal was separated by a beam splitter (560/10 nm) and simultaneously detected with a band-pass filter for SR101 (610/75 nm) and GCaMP6 (525/50 nm). A single image plane consisted of 512 × 512 pixels, and in-plane pixel size was 0.9 μm × 0.9 μm. The methods for functional imaging using two-photon microscopy have been reported in detail^38^. Briefly, continuous image capturing was conducted for neurons on the surface over the barrel cortex at a rate of 0.25 s per frame for 60 s with a 512 × 512 pixel field of view. Percentage change in green fluorescence of neurons was manually measured offline with LAS AF software (Leica Microsystems GmbH). Sensory-evoked neuronal excitation was examined by means of whisker stimulation as reported previously^38^. Air puff stimulations (15 psi, 50 ms pulse width, 10 s duration, 10 Hz) were given to whiskers on the contralateral side of scanning 4 times in a day.

### Electrophysiology

Electrophysiological recordings were performed on one monkey while seated in a monkey-chair with its head fixed via a headpost, and maintained under an anesthetized condition with intravenous infusion of propofol (20 mg/kg/h). Local field potential (LFP) was recorded using a multi-contact linear probe with twelve recording contacts separated by 150 μm (Axial Array, FHC, Bowdoin, USA). The probe was inserted into the brain with a hydraulic micromanipulator (MO-97A, Narishige, Tokyo, Japan) guided with a stainless steel tube and an implanted recording chamber system (Crist Instruments, Hagerstown, USA). Chamber grids were either purchased from Crist Instruments or custom-made with a 3D printer (Object260 Connex3, Stratasys, Eden Prairie, USA). For every recording session, the monkey was CT-scanned before the recording probe was removed. Electrode location relative to the hM_3_Dq expression was confirmed by aligning the CT image with the pre-acquired PET image on PMOD®. The probe was connected to a multichannel acquisition system (System 3, Tucker-Davis Technologies (TDT), Alachua, USA) running on a Windows PC. For LFP, the signal was band-pass filtered between 1.5 Hz and 500 Hz, and sampled at 3 kHz. LFP signals were put to further analysis only when multi-unit waveforms were qualitatively observed in a high frequency band (i.e., 400 Hz – 5 kHz) at the corresponding recording sites. The spectral component of the LFP signal was analyzed with MATLAB (MathWorks, Natick, USA). Spectral power of each frequency was computed using a short FFT algorithm within a 60-s window sliding in 30-s steps. Change in the gamma band power (40 Hz) and its change relative to that in a 3- or 10-min pre-injection period was quantified. The spectrogram was smoothed with a Hanning window with 4.3-min half-maximum width for display reasons.

### Behavioral testing

Three monkeys (#226, #229, #245) were tested with a spatial delayed response task (Fig. 7c). The protocol was based on previous studies using the Wisconsin general testing apparatus^39,40^. Behavioral testing was conducted in a sound-attenuated room. Monkeys were seated in a monkey chair from which they could reach out one hand and take food to their mouths. A wooden table with two food-wells was placed in front of the monkeys, and a screen was placed between the monkeys and the table. First, a piece of food reward (raisin or half peanut) was placed in one of the two food-wells, and then both wells were covered with wooden plates. Then, the screen was placed for 0.5, 5 or 10 s, which served as delay periods (0.5-s delay was not introduced for non-DREADD monkeys). The position of the baited well (left or right) was determined pseudo-randomly. After the delay period, the screen was removed and the monkeys were allowed to select either food-well to get the food. The monkeys were allowed to take the food if they reached for the correct food-well and removed the cover plate. The inter-trial interval was set at 10-15 s. A daily session lasted about one hour, and consisted of 3-4 blocks of 20-30 trials, which were interleaved by a 5-min rest period. The behavioral testing began immediately or 1 h after an i.m. administration of either vehicle (2% DMSO in saline) or DCZ (100 μg/kg), and was also conducted on the next day (24 h later) of the DCZ administration. One of the monkeys injected with AAV-hM_4_Di in the bilateral PFC (#229) was also tested in a non-memory control task, which was almost the same as the delayed response task except that the screen was not placed during the delay period and that the 0.5-s delay was not introduced.

Three monkeys without AAV injections (non-DREADD; #224, #228, #230) were tested with a reward-size task (Supplementary Fig. 6a) using the same protocol as applied in a previous study^41^. The behavioral testing began 10 min after an i.m. administration of either vehicle (2% DMSO in saline) or DCZ (100 μg/kg).

### Histology and immunostaining

The monkeys were deeply anesthetized with an overdose of sodium pentobarbital (80 mg/kg, i.v.) and transcardially perfused with saline at 4°C, followed by 4% paraformaldehyde in 0.1 M phosphate buffered saline (PBS), pH 7.4. The brains were removed from the skull, postfixed in the same fresh fixative overnight, saturated with 30% sucrose in phosphate buffer (PB) at 4°C, and then cut serially into 50-μm-thick sections on a freezing microtome. For visualization of immunoreactive signals of hM_4_Di, a series of every 6th section was immersed in 1% skim milk for 1 h at room temperature and incubated overnight at 4°C with rabbit anti-M_4_ polyclonal antibody (1:200; H-175, Millipore, Burlington, USA) in PBS containing 0.1% Triton X-100 and 1% normal donkey serum for 2 days at 4°C. The sections were then incubated in the same fresh medium containing biotinylated donkey anti-rabbit IgG antibody (1:1,000; Jackson ImmunoResearch, West Grove, USA) for 2 h at room temperature, followed by avidin-biotin-peroxidase complex (ABC Elite, Vector Laboratories, Burlingame, USA) for 2 h at room temperature. For visualization of the antigen, the sections were reacted in 0.05 M Tris-HCl buffer (pH 7.6) containing 0.04% diaminobenzidine (DAB), 0.04% NiCl2, and 0.003% H_2_O_2_. The sections were mounted on gelatin-coated glass slides, air-dried, and cover-slipped. A part of other sections was Nissl-stained with 1% Cresyl violet. The same protocol was used for visualization of immunoreactive signals of hM_3_Dq with rabbit anti-M_3_ polyclonal antibody (1:200; HPA024106, Atlas Antibodies, Stockholm, Sweden).

The mice were deeply anesthetized with sodium pentobarbital and then transcardially perfused with PBS. Brain tissues were removed and fixed with 4% paraformaldehyde in PB overnight, followed by cryoprotection with 30% sucrose in PB. 10-μm-thick frozen sections were generated in a cryostat (HM560; Carl Zeiss, Oberkochen, Germany). Visualization of immunoreactive signals of hM_4_Di was performed using an anti-M_4_ antibody (H-175) and fluorophore-conjugated secondary antibodies (Invitrogen, Carlsbad, USA) as described previously^15^.

Images of sections were digitally captured using an optical microscope equipped with a high-grade charge-coupled device (CCD) camera (Biorevo, Keyence, Osaka, Japan).

### Pharmacokinetics analysis

Four monkeys and 15 mice were used to assess the concentration of DCZ and its metabolites in plasma, cerebrospinal fluid (CSF) or brain. In monkeys, blood and CSF were collected at 15, 30, 45, 60, 75, 90, 105, and 120 min after DCZ administration (100 μg/kg, i.v. or i.m.) under ketamine and xylazine anesthesia. In mice, after intraperitoneal injection of DCZ (100 μg/kg), blood and brain samples were collected at 5, 30, 60, 120 min or 24 h, and CSF was collected at 30 and 120 min. Mouse blood was collected by heparinized syringes via the heart under isoflurane anesthesia followed by centrifugation at 10,000 g for 5 min to obtain plasma samples, and brains were removed and frozen by liquid nitrogen immediately after blood collection. All samples were stocked at −80°C until analysis.

The protocols for sample pretreatment of CSF and plasma were described previously^6^. Mouse brains were homogenized in 2-brain-weight water. An acetonitrile solution (0.8 mL) containing granisetron (0.5 ng/mL) as internal standard was added to the brain homogenate (0.2 mL), followed by centrifugation at 10,600 g for 2 min at 4°C. The supernatant was collected and filtered by solid phase extraction column (Phree™; Shimadzu GLC Ltd, Tokyo, Japan). The filtrate was dried under nitrogen gas at 40°C and re-dissolved in 5% acetonitrile (0.2 mL) followed by sonication for 30 s and centrifugation (500 g) for 2 min. The supernatant was filtered by solid phase extraction column again before its application to LC/MS/MS.

Quantification of C21, DCZ, and DCZ-N-oxide was performed by multiple reaction monitoring (MRM) using a Shimadzu UHPLC LC-30AD system (Shimadzu Corp., Kyoto, Japan) coupled to a tandem MS AB Sciex Qtrap 6500 system (AB Sciex LLC, Framingham, USA). The following MRM transitions (Q1/Q3) were used to monitor each compound: C21 (279.0/193.0), DCZ (293.0/236.0), DCZ-N-oxide (309.0/192.9), granisetron (313.2/138.1). Other protocols were described previously^6^.

### Statistical analyses

#### In vitro assays

All data were analyzed using GraphPad Prism 7 (San Diego, USA). Inhibition binding data were analyzed according to a one-site binding model, and the equilibrium dissociation constants (*Ki*) of unlabeled ligands were calculated by constraining the radioligand *Kd* to the values estimated from saturation binding assays. Concentration−response curves were fitted to a three-parameter logistic equation. All affinity and potency values were estimated as logarithms.

#### PET imaging

To estimate the specific binding of [^11^C]DCZ, regional binding potential relative to nondisplaceable radioligand (BP_ND_) was calculated by PMOD® with an original multilinear reference tissue model (MRTMo)^42^ as described by the following equation:

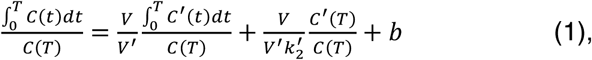

where C(t) and C’(t) are the regional or voxel time-radioactivity concentrations in the tissue and reference regions, respectively (kBq/mL), V and V’ are the corresponding total distribution volumes (mL/mL), k_2_’ (min^−1^) is the clearance rate constant from the reference region to plasma, and b is the intercept term, which becomes the constant for T > t*. In this study, t* was determined as 2 min for mice studies and 15 min for monkey studies. Eq. 1 allows estimation of three parameters, β_1_ = V/V’, β_2_ = V/(V’k_2_’), and β_3_ = b by multilinear regression analysis for T > t*. Assuming that the nondisplaceable distribution volumes in tissue and reference regions are identical, BP_ND_ (BP_ND_ = V/V’ – 1) is calculated from the first regression coefficient as BP_ND_ = (β_1_ – 1).

Estimates of fractional occupancy were calculated by the following equation:

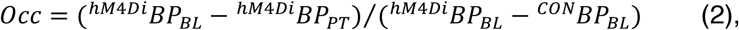

where ^*hM4Di*^*BP*_*BL*_ and ^*CON*^*BP*_*BL*_ indicate BP_ND_ at the hM_4_Di-expressing putaminal region and control putamen under baseline conditions, respectively, while ^*hM4Di*^*BP*_*PT*_ indicates BP_ND_ at the hM_4_Di-expressing putaminal region under pretreatment condition. The relationship between occupancy (*Occ*) and agonist dose (*D*_*agonist*_) was modeled by the Hill equation

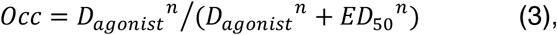

where *ED*_*50*_ and *n* indicate the agonist dose achieving 50% occupancy and the Hill coefficient, respectively.

For FDG-PET analysis, dynamic SUV images were motion-corrected and then were averaged between 30-60 min after injection of the radioligand. Voxel-wise statistical analyses were performed with SPM12 software (Wellcome Department of Cognitive Neurology, London, UK; www.fil.ion.ucl.ac.uk) and MATLAB R2016a (MathWorks Inc., Natick, USA). We used a total of 12 scans including vehicle and DCZ 3 μg/kg pretreatment conditions to detect the metabolic change by hM_3_Dq activation and a total of 12 scans including vehicle and DCZ 100 μg/kg pretreatment conditions to examine the effect of off-target actions in the non-DREADD monkeys. The averaged SUV images were spatially normalized into a standard template brain MR image^43^ after coregistration of individual MR images. The resulting images were smoothed with a 2.6-mm Gaussian filter and then extracted by brain mask. The SUV values for each scan were corrected using grand mean scaling and by analysis of covariance for global normalization. Subsequently, repeated measures one-way ANOVA was applied. The statistical threshold was set at uncorrected *p* < 0.001 (T > 4.50) and an extent of 100 contiguous voxels per cluster. In VOI-based analysis, we used a total of 17 scans including vehicle and a DCZ dose of 1 μg/kg, 3 μg/kg pretreatment conditions obtained from hM_3_Dq-expressing monkeys and a total of 16 scans including vehicle, and a DCZ dose of 1 μg/kg, 100 μg/kg pretreatment conditions obtained from non-DREADD monkeys. The SUV values were obtained by PMOD® and then were normalized by whole-brain value. Paired t-test was used to detect the metabolic change by hM_3_Dq activation in each condition. To examine the dose-dependency or off-target actions, one-way ANOVA followed by post-hoc Tukey-Kramer test was performed.

#### Neuronal activity

Time-dependent changes in fluorescent signal and LFP power were analyzed according to a one-phase exponential association model using GraphPad Prism 7. Repeated measures ANOVA followed by post-hoc Dunnett test was used to examine differences between baseline and drug-evoked mean neural signals. The signals were displayed after normalization to the baseline levels, but statistical tests were conducted on the original data.

To examine the long-term effect of hM_3_Dq activation on normal neuronal responsiveness, fluorescent signal change within 0.2 – 10.2 s after whisker stimulation was averaged across 4 trials and compared using pair-wise t-test. DCZ-induced fluorescent signal changes at 10 min after the 1st and 2nd DCZ administrations were compared by pair-wise t-test. The second DCZ administration was applied 24 h after the first.

#### Behavioral testing

To examine the effect of DCZ on the performance of the delayed response task, behavioral measurement (correct rates) was subjected to two-way ANOVA (treatment × delay) and the following post-hoc t-test with Bonferroni correction using GraphPad Prism 7. To examine whether the performance was recovered at 24 h after DCZ administration, the same procedures were conducted between DCZ and post-DCZ, and vehicle and post-DCZ sessions. To examine the effect of DCZ treatment on the behavior in test time, the data of 5- and 10-s delays were pooled and then compared between treatment conditions by t-test. To examine the effect of DCZ on the performance of the reward-size task, reaction times and error rates were subjected to two-way ANOVA (treatment × reward size). The total correct trial number for each session was subjected to two-way ANOVA (treatment × subject). For each analysis, effect size eta-squared (η^2^) was calculated using R.

#### General data collection and statistical statements

Data collection and analyses were not performed blinded as to the conditions of the experiments, except in vitro studies which were blinded. For *in vivo* experiments, the order of drug administration (e.g., DCZ or vehicle) was pseudo-random and shuffled across animals. No samples were excluded from the analysis. Statistical tests included parametric and nonparametric methods. All statistical tests were two-sided unless specifically stated otherwise. In some statistical tests, data distributions were assumed to be normal and/or with equal variances, but this was not formally tested. No statistical methods were used to predetermine sample sizes, but our sample sizes were similar to those reported in previous publications (for example, see refs^5-8,10,30^).

#### Reporting Summary

Further information on research design is available in the Nature Research Reporting Summary linked to this article.

## Data availability

The data that support the findings of this study are available from the corresponding author upon reasonable request

## Code availability

The code to generate the results and the figures of this study are available from the corresponding author upon reasonable request.

