## Supplementary Figs 1-6 and Tables 1-2 for "Deschloroclozapine, a potent and selective chemogenetic actuator enables rapid neuronal and behavioral modulations in mice and monkeys"

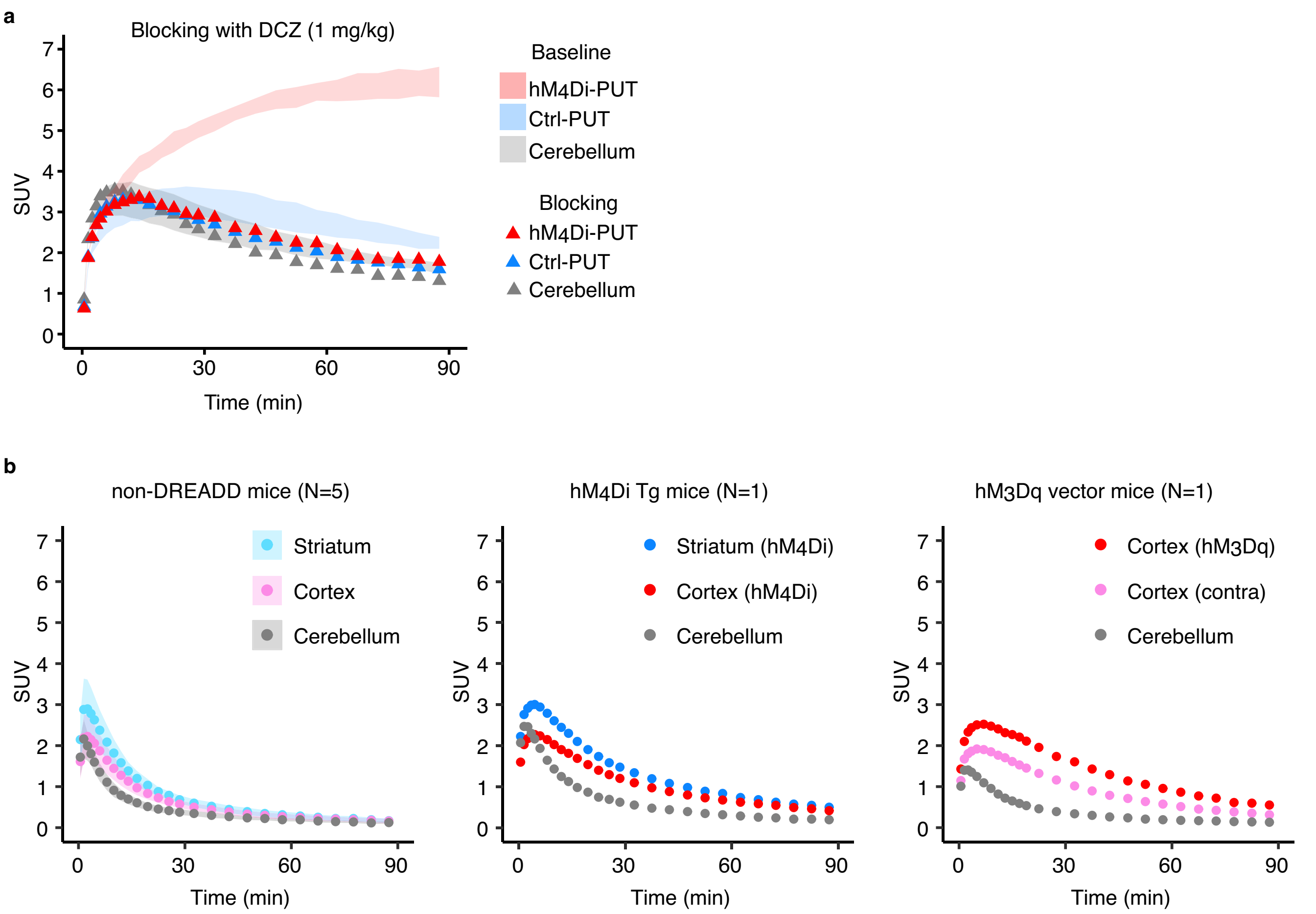


**Supplementary Figure 1**

**Blocking with non-labeled DCZ.**

**a,** Time course of regional uptake of $[^{11}C]$DCZ under blocking with non-radiolabeled DCZ (1 mg/kg, i.v.) at the putative hM$_4$Di-expressing region (hM$_4$Di-PUT), contralateral control region (Ctrl-PUT), and the cerebellum in monkey #209. Shaded areas represent baseline uptakes (mean ± sd, n = 3; same as those shown in Figure 1g). **b,** *Left*: Time course of regional uptake of $[^{11}C]$DCZ (mean ± sd) at the striatum (blue), cortex (red), and cerebellum (gray) in 5 wild-type mice. *Center* and *Right*: Time course of regional uptake of $[^{11}C]$DCZ in an hM$_4$Di-Tg (center) and a mouse expressing hM$_3$Dq in the barrel cortex (right, cf. Fig. 4d), respectively. SUV indicates standardized uptake value [regional radioactivity (Bq/cm$^3$) × body weight (g) / injected radioactivity (Bq)].

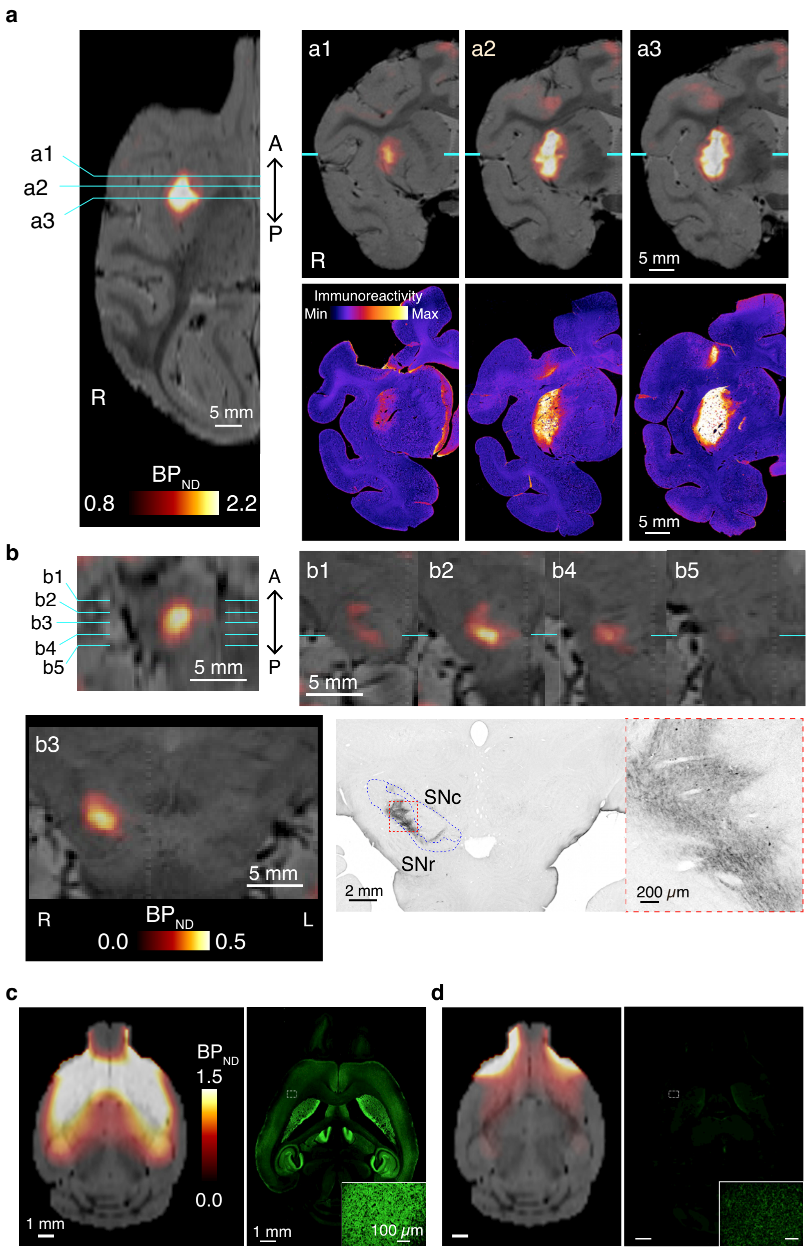


**Supplementary Figure 2**

**[$^{11}$C]DCZ-PET visualizes hM$_4$Di expression in monkeys and mice.**

**a,** Horizontal (top left) and coronal sections of parametric image of specific binding of [$^{11}$C]DCZ overlaying MR image (a1-a3) and corresponding anti-hM$_4$ staining sections (bottom) including putamen in monkey #209. Relative density of staining is color-coded (see scale). **b,** Horizontal (top left) and coronal sections of parametric image of specific binding of [$^{11}$C]DCZ overlaying MR image (B1-B5) and corresponding anti-hM$_4$ staining sections (bottom right) including substantia nigra in monkey #209. A high-magnification image of the red boxed area is shown in the right panel focusing on immuno-positive axons in substantia nigra pars reticulata (SNr). We obtained similar observations from another monkey (#212). **c,** Horizontal section of parametric image of specific binding of [$^{11}$C]DCZ overlaying MR image (left), immunohistochemical image with an anti-hM$_4$ antibody of the corresponding horizontal section (right), and a high magnification image (inset) of the boxed area of a hM$_4$Di-transgenic mouse. **d,** Same as **c** but for a wild-type mouse.

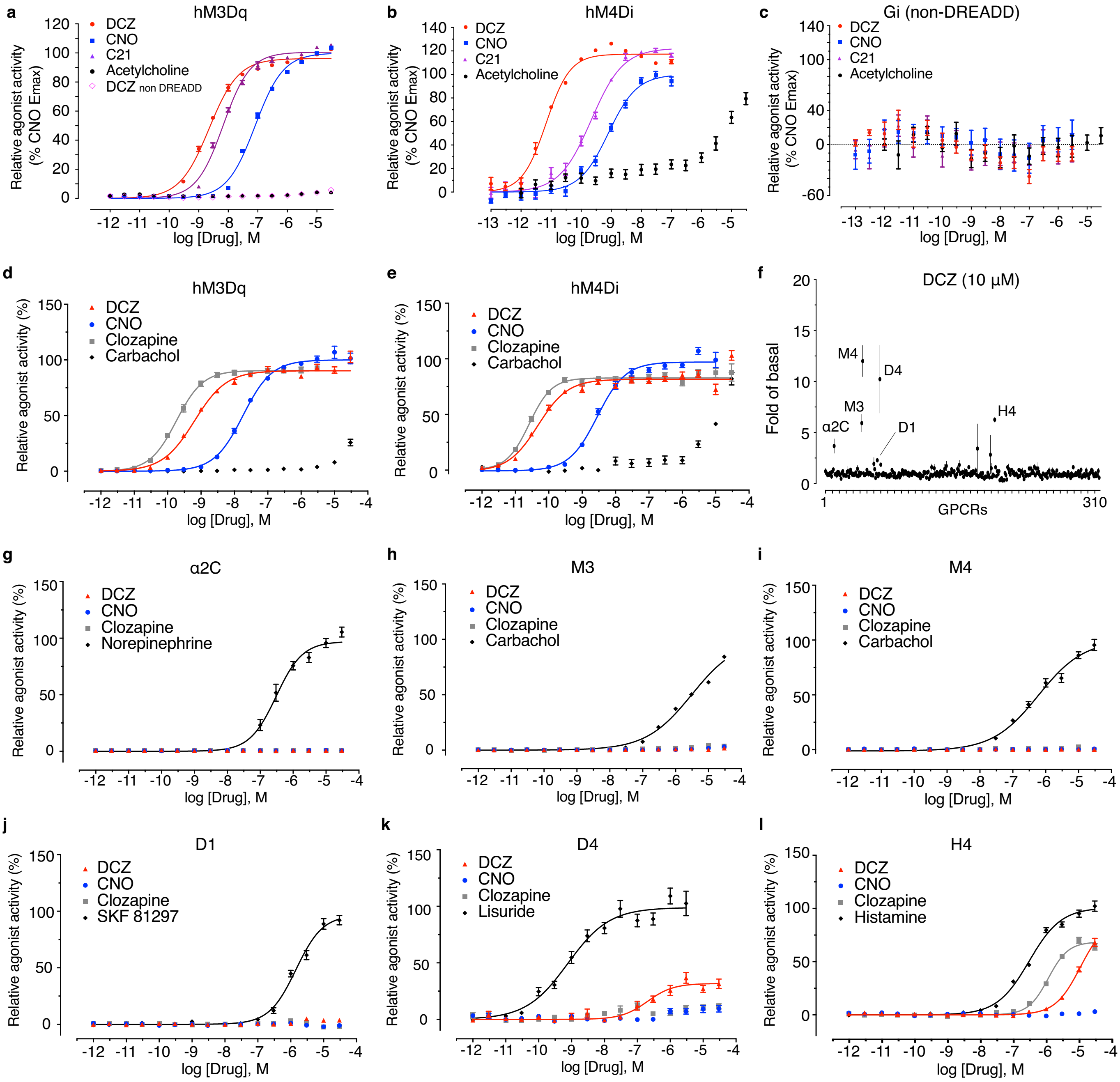


**Supplementary Figure 3**

**Selective agonist potency of DCZ on DREADDs *in vitro*.**

**a,** Concentration-response effect of DCZ, CNO, C21 and acetylcholine on $Ca^{2+}$ mobilization in $hM_3Dq$- expressing HEK293T cells, respectively; DCZ, ${}^{hM3Dq}EC_{50}$ = 2.3 (2.1-2.5) nM, CNO [${}^{hM3Dq}EC_{50}$ = 82 (76-89) nM], C21 [${}^{hM3Dq}EC_{50}$ = 6.7 (6.2-7.4) nM]. DCZ $_{non\text{-}DREADD}$ indicates effect of DCZ on HEK293T cells not expressing DREADDs (pcDNA-transfected cells). **b,** Concentration-response effect of DCZ, CNO, C21 and acetylcholine on cAMP inhibition in $hM_4Di$-expressing HEK293T cells, respectively; DCZ, ${}^{hM4Di}EC_{50}$ = 0.0048 (0.0032-0.0071) nM, CNO [${}^{hM4Di}EC_{50}$ = 0.49 (0.37-0.65) nM], C21 [${}^{hM4Di}EC_{50}$ = 0.15 (-0.12-0.19) nM]. **c,** Same as **b** but for the use of pcDNA-transfected cells. **d,e,** Concentration-response effect of DCZ, CNO, clozapine and carbachol on β-arrestin recruitment of $hM_3Dq$ and $hM_4Di$, respectively; DCZ, ${}^{hM3Dq}EC_{50}$ = 0.63 nM, ${}^{hM4Di}EC_{50}$ = 0.048 nM; clozapine, ${}^{hM3Dq}EC_{50}$ = 0.18 nM, ${}^{hM4Di}EC_{50}$ = 0.026 nM; CNO, ${}^{hM3Dq}EC_{50}$ = 18 nM, ${}^{hM4Di}EC_{50}$ = 2.8 nM. For panels **a-e**

data represent mean ± sem of N=3 biological replicates. **f,** DCZ was screened against 318 GPCRs for antagonism in β-arrestin recruitment assay. Each point indicates normalized activity at 10 μM to the baseline level at a given GPCR for N=4 technical replicates. Although DCZ increased β-arrestin recruitment >2-fold of baseline at several GPCRs, including adrenaline $\alpha_{2C}$, muscarinic acetylcholine $M_3$ and $M_4$, dopamine $D_1$ and $D_5$, and histamine $H_4$ receptors, follow-up concentration response studies (shown in **g-l**) revealed that DCZ exhibited minimal activity on these receptors ($EC_{50}$ > 1 μM). **g-l,** Concentration-response effect of DCZ, CNO, clozapine and control agonist on β-arrestin recruitment of 6 target receptors for a single experiment which was replicated twice with mean ± sem of triplicate determinations: $\alpha_{2C}$ **(g),** $M_3$ **(h),** $M_4$ **(i),** $D_1$ **(j),** $D_5$ **(k),** $H_4$ **(l)**. Data are shown as mean ± sem. Note that the expression levels of $M_3$ and $M_4$ were similar to their corresponding DREADDs; $M_3$ [372 (238 - 458) fmol/mg]; $M_4$ [190 (120 - 348) fmol/mg; hM3Dq [597 (242 - 846) fmol/mg]; h$M_4$Di [238 (154 - 328) fmol/mg]; mean (range), respectively.

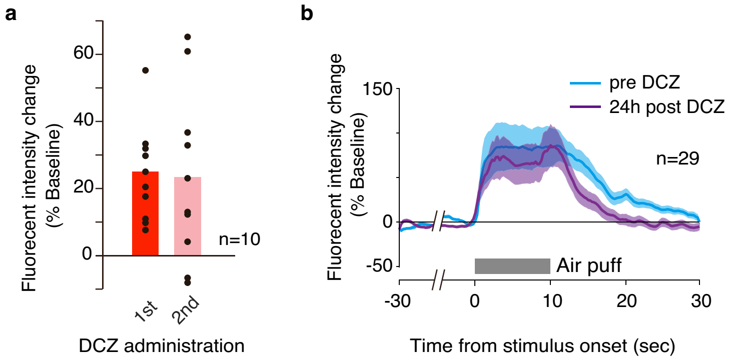


**Supplementary Figure 4**

**Safety of chemogenetic activation with DCZ *in vivo*.**

**a,** Comparison of chemogenetically induced changes in fluorescence signals between 1st and 2nd DCZ administration (100 µg/kg, i.p.), separated by 24 hours. Data were collected 10 min after DCZ administrations. No significant difference was found ($p = 0.83$; paired t-test, n=10 cells). **b,** Comparison of stimulus-evoked response of fluorescence signals before (pre-DCZ, cyan) and 24 h after DCZ administration (purple). Curves and shaded areas represent mean and sem. No significant difference was found between the two conditions ($p = 0.53$; paired t-test, n=29 cells). Data were collected from one animal.

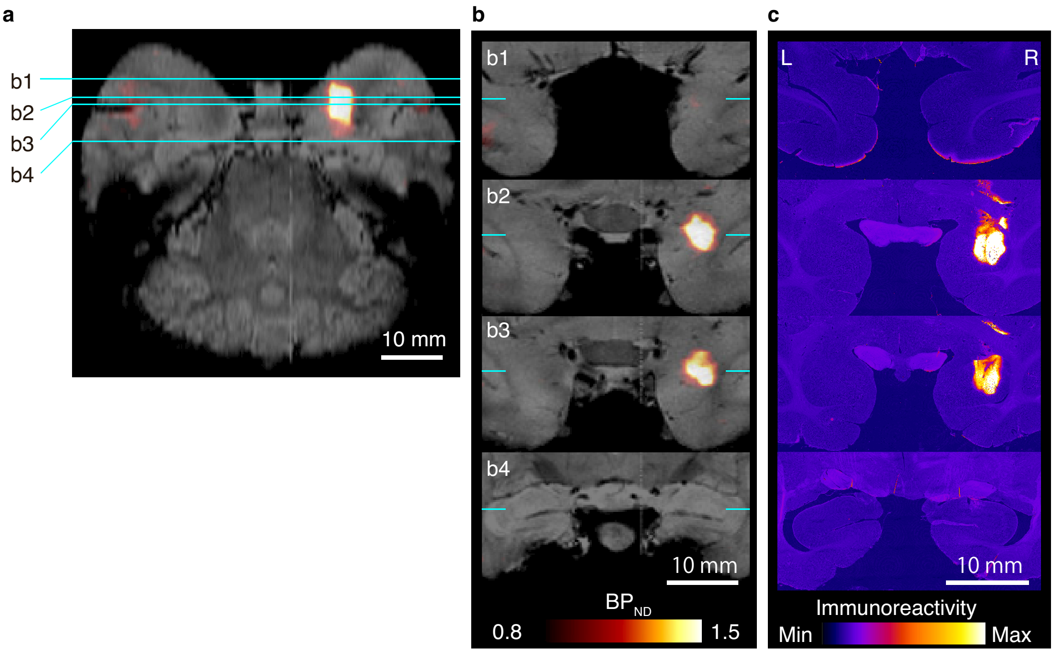


**Supplementary Figure 5**

**[11C]DCZ-PET visualizes hM₃Dq expression in monkeys.**

**a,b,** Horizontal and coronal sections of parametric image of specific binding of [$^{11}$C]DCZ overlaying MR image in monkey #233, respectively. The positions of sections b1-b4 are indicated by cyan lines in **a**. **c,** Anti-hM$_3$ stained sections corresponding to b1-b4. Relative density of staining is color-coded (see scale). We obtained similar observations from another monkey (#214).

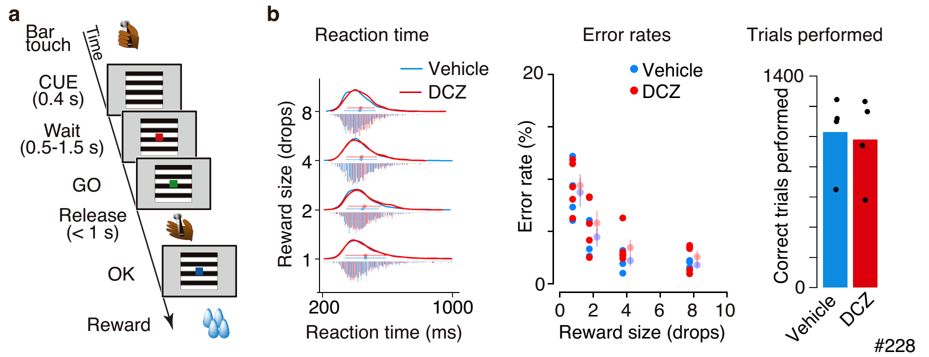


**Supplementary Figure 6**

**Negative behavioral control.**

**a,** Illustration of reward-size task. Each trial in this task began when the monkey touched a lever, which was followed by the appearance of a visual cue signaling the size of the upcoming reward (1, 2, 4 or 8 drops). To obtain a reward, the monkeys had to release the lever when a visual target changed color from red to green. **b,** Example behavioral data of reward size task are shown for DCZ (100 µg/kg, i.m.; red; total 4,730 trials/4 sessions) and vehicle administration session (cyan; 5,109 trials/4 sessions) obtained from a non-DREADD monkey (#228). *Left:* Distribution of reaction time for each reward size. No main effect of treatment, two-way ANOVA, treatment × reward size, effect size $\eta^2 = 0.0007$, 0.016, 0.012 for three monkeys, respectively. *Middle:* Error rates as a function of reward size. Left dots represent single session data while right dots and vertical bars indicate mean ± sem. No main effect of treatment, $\eta^2 = 0.012$, 0.04, and 0.012, respectively. *Right:* Number of correct trials performed. No effect of treatment, two-way ANOVA, treatment × subject, $\eta^2 = 0.0001$ (data from three monkeys were pooled because of small sample size).

**Supplementary Table 1**

**Binding affinities of DCZ and clozapine to endogenous receptors, channels and transporters.**

| | $K_i$ *(nM)* | |
|---|---|---|
| | DCZ | clozapine |
| 5-HT$_{1A}$ | 410 | 110 |
| 5-HT$_{1B}$ | >10,000 | 400 |
| 5-HT$_{1D}$ | 2,800 | 2,100 |
| 5-HT$_{1E}$ | 3,400 | 970 |
| 5-HT$_{2A}$ | 87 | 13 |
| 5-HT$_{2B}$ | 100 | 4.5 |
| 5-HT$_{2C}$ | 320 | 29 |
| 5-HT$_3$ | 110 | 240 |
| 5-HT$_4$ | >10,000 | 3,900 |
| 5-HT$_{5A}$ | 4,500 | 17 |
| 5-HT$_6$ | 640 | 18 |
| 5-HT$_7$ | 190 | 12 |
| $\alpha_{1A}$ | 2,800 | 1.6 |
| $\alpha_{1B}$ | >10,000 | 7 |
| $\alpha_{1D}$ | >10,000 | 350 |
| $\alpha_{2A}$ | 780 | 140 |
| $\alpha_{2B}$ | 500 | 27 |
| $\alpha_{2C}$ | >10,000 | 34 |
| $\beta_1$ | >10,000 | 10,000 |
| $\beta_2$ | >10,000 | 10,000 |
| $\beta_3$ | >10,000 | 10,000 |
| BZP | >10,000 | 10,000 |
| $D_1$ | 510 | 190 |

| | $K_i$ *(nM)* | |
|---|---|---|
| | DCZ | clozapine |
| $D_2$ | >10,000 | 430 |
| $D_3$ | >10,000 | 650 |
| $D_4$ | 1,000 | 39 |
| $D_5$ | 1,800 | 240 |
| DAT | >10,000 | >10,000 |
| DOR | >10,000 | 6,800 |
| GABA$_A$ | >10,000 | >10,000 |
| $H_1$ | >10,000 | 2 |
| $H_2$ | 1,100 | 150 |
| $H_3$ | 4,600 | >10,000 |
| $H_4$ | 1,100 | 820 |
| $M_1$ | 83 | 14 |
| $M_2$ | 320 | 14 |
| $M_3$ | 230 | 21 |
| $M_4$ | 210 | 11 |
| $M_5$ | 55 | 94 |
| MOR | >10,000 | >10,000 |
| NET | 3,500 | 3,200 |
| PBR | >10,000 | >10,000 |
| SERT | >10,000 | 1,600 |
| $\sigma_1$ | >10,000 | >10,000 |
| $\sigma_2$ | >10,000 | 1,600 |

*Ki* values are the average of at least 3 triplicate experiments with standard deviation values 3-fold less than the average. BZP, benzodiazepine receptor; DAT, dopamine transporter; DOR, δ-opioid receptor; MOR, μ-opioid receptor; NET, norepinephrine transporter; PBR, peripheral benzodiazepine receptor; SERT, serotonin transporter.

**Supplementary Table 2**

**Summary of monkeys used in the study.**

| ID | Species | Sex | Age (y) | Vector injection | DCZ-PET | FDG-PET | Histology | PK | Rec. | Behavior |
|---|---|---|---|---|---|---|---|---|---|---|
| 176 | J | M | 10 | | | | | ✔ | | |
| 209 | J | M | 4 | hM4Di putamen | ✔ | | ✔ | | | |
| 210 | J | M | 5 | | | ✔ | | | | |
| 212 | R | F | 9 | hM4Di putamen | ✔ | | | | | |
| 214 | R | F | 8 | hM3Dq amygdala | ✔ | ✔ | | | | |
| 215 | R | F | 9 | hM3Dq amygdala | ✔ | ✔ | | | | |
| 224 | J | M | 8 | | | | | | | ✔ |
| 226 | J | F | 7 | | | | | | | ✔ |
| 227 | J | M | 8 | | | | | ✔ | | |
| 228 | R | M | 5 | | | | | | | ✔ |
| 229 | R | M | 5 | hM4Di PFC | ✔ | | | | | ✔ |
| 230 | R | M | 4 | | | | | ✔ | | ✔ |
| 231 | R | M | 4 | | | ✔ | | ✔ | | |
| 233 | J | M | 4 | hM3Dq amygdala | ✔ | | ✔ | | ✔ | |
| 236 | J | M | 5 | hM3Dq amygdala | ✔ | ✔ | | | | |
| 245 | J | F | 7 | hM4Di PFC | ✔ | | | | | ✔ |
| Total | 16 | | | 8 | 8 | 5 | 2 | 4 | 1 | 6 |

A tick indicates the subject was used in the experiment. Species (J, Japanese; R, Rhesus), Sex (F, Female; M, Male), Age (years), PK, Pharmacokinetics study, Rec., Electrophysiological recording study.
